# Promoting Polarization and Differentiation of Primary Human Salivary Gland Stem/Progenitor Cells in Protease-Degradable Hydrogels via ROCK Inhibition

**DOI:** 10.1101/2024.11.24.625065

**Authors:** Apoorva S. Metkari, Robert L. Witt, David M. Cognetti, Charles Dhong, Xinqiao Jia

## Abstract

Towards the goal of *in vitro* engineering of functional salivary gland tissues, we cultured primary human salivary stem/progenitor cells (hS/PCs) in hyaluronic acid-based matrices with varying percentages of proteolytically degradable crosslinks in the presence of Rho kinase (ROCK) inhibitor. Single cells encapsulated in the hydrogel grew into organized multicellular structures by day 15, and over 60% of the structures developed in the non-degradable and 50% degradable hydrogels contained a central lumen. Importantly, ROCK inhibition led to the establishment of multicellular structures that were correctly polarized, as evidenced by apical localization of a Golgi marker GM130, apical/lateral localization of tight junction protein zonula occludens-1 (ZO-1), and basal localization of integrin β1 and basement membrane proteins laminin α1 and collagen IV. Cultures maintained in 50% degradable gels with ROCK inhibition exhibited an increased expression of acinar markers *AQP5* and *SLC12A2* (at the transcript level) and AQP5 and NKCC1 (at the protein level) as compared to those without ROCK inhibition. Upon stimulation with isoproterenol, α-amylase secretion into the lumen was observed. Particle-tracking microrheology was employed to analyze the stiffness of cells using mitochondria as the passive tracer particles. Our results indicated that cells grown in 100% degradable gels were stiffer than those maintained in non-degradable gels, and cells cultured with the ROCK inhibitor were softer than those maintained without the inhibitor. We conclude that reducing cellular contractility via ROCK inhibition while retaining some degree of matrix confinement promotes the establishment of multicellular structures containing pro-acinar cells with correct apicobasal polarization.

## INTRODUCTION

Human salivary glands produce and secrete saliva through coordinated actions of acinar, myoepithelial, and ductal cells. Each gland contains secretory end buds connected to branched ducts for efficient production and transport of saliva.^1^ Standard radiation therapy for head and neck malignancies causes irreversible loss of saliva-producing acinar cells, giving rise to xerostomia that is manifested as hyposalivation.^2, 3^ Due to reduced saliva flow, patients have higher rates of dental caries, periodontal diseases, and oral infections, as well as reduced overall health, nutrition, and quality of life.^4^ Current clinical remedies for xerostomia aim to protect the salivary gland during radiation treatment, palliate symptoms using water, oral sialagogues, or anti-inflammatory agents, or stimulate the secretory function using cholinergic muscarinic receptor agonists. None has provided long-term therapeutic benefits.^5^ Engineering of implantable and functional salivary gland mimetics will offer a potential long-term curative solution to xerostomia.

*In vitro* engineering of functional salivary glands requires reprogrammable autologous progenitor cells, soluble instructive signals, and permissive matrices to aid coordinated tissue growth. We have established methods for isolating, expanding, and differentiating adult human salivary stem/progenitor cells (hS/PCs).^6^ Our customized, hyaluronic acid (HA)-based hydrogels are proven to facilitate the development of multicellular spheroids from dispersed hS/PCs. We identified crosslinking chemistry, gel stiffness, and matrix degradability (Figure 1A) that promote the rapid development of pro-acinar spheroids.^7–11^ However, a cohesive and mechanically robust basement membrane was not established around the developing microstructures. Although occasionally structures with a lumen were seen, under neurotransmitter stimulation, the multicellular structures secreted α-amylase outwards into the hydrogel but not into the lumen, suggesting that cells are not correctly polarized and/or the structures are not sealed.^12^

**Figure 1.**
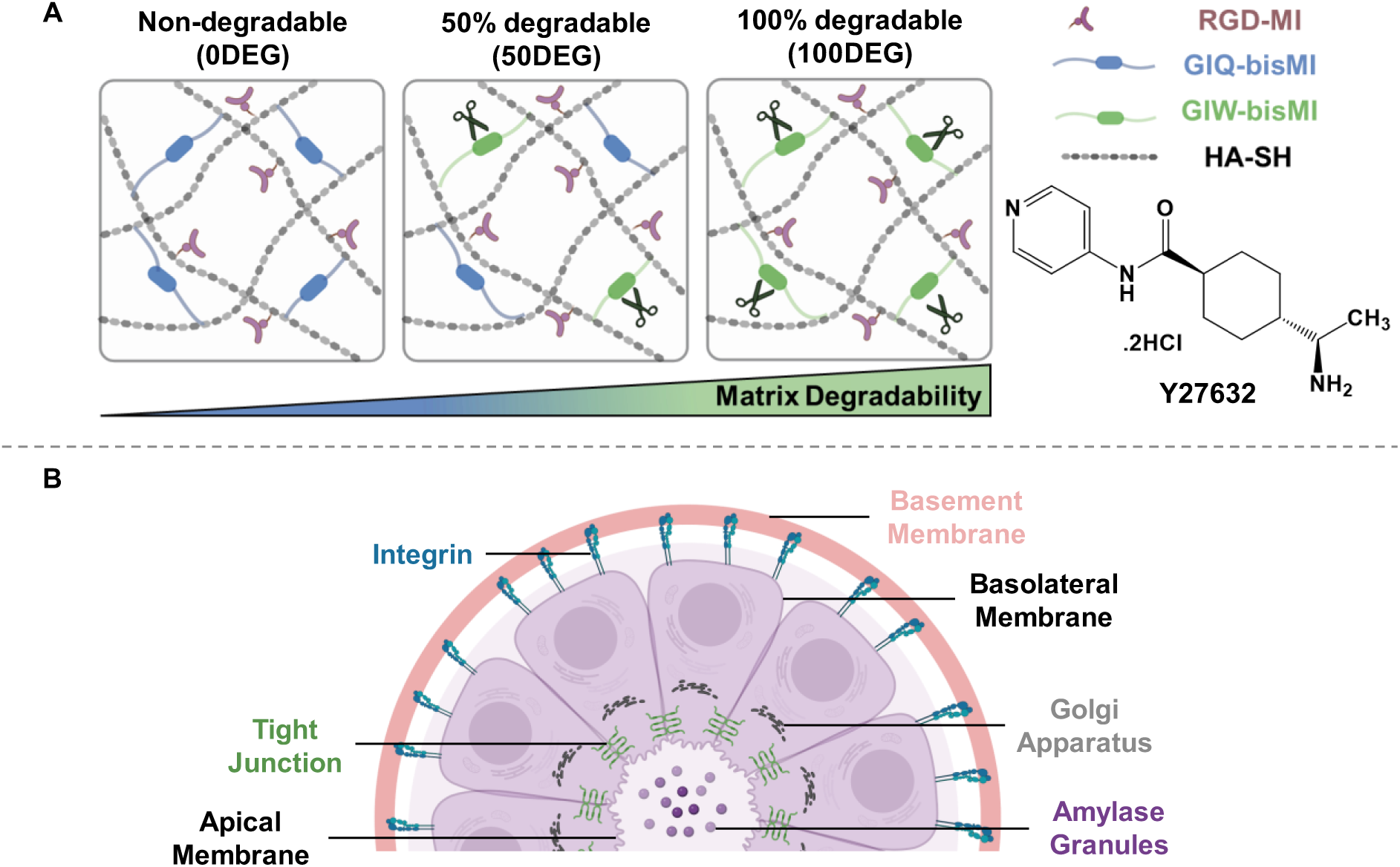
Development of polarized hS/PC spheroids in customized HA gels in the presence of a ROCK inhibitor Y27632. **(A)** Hydrogels were prepared with the same cell adhesive ligand (RGD-MI) and crosslinker (bisMI) concentrations but varying ratios of MMP-degradable (GIW) and non-degradable (GIQ) crosslinkers. hS/PCs were encapsulated in the hydrogel as single cells, and media were supplemented with ROCK inhibitor Y27632 to stimulate lumen formation. **(B)** Schematic depicting spatial localization of proteins in correctly polarized epithelial structures. The apical membrane faces the lumen, and the basal membrane is in contact with the basement membrane. Golgi is in between the nucleus and the apical membrane. Tight junction proteins are located apically and laterally. Integrins are located basally to direct cell attachment to the basement membrane. Amylase is secreted directionally into the lumen.

During the development of mammalian epithelial organs, a lumen can form through selective apoptosis of cells not in contact with the extracellular matrix (ECM). Alternatively, a space between two closely apposed cells can be created *de novo* by trafficking vesicles containing the apical membrane to a space between cells or in a single cell.^13^ A pre-requisite for lumen formation is the acquisition of collective apicobasal polarity.^14–16^ Attainment of proper polarity requires cells to perceive and integrate polarizing cues from the cellular and extracellular milieu. A cohesive ECM, specifically the basement membrane, provides the initial cue for orienting the apicobasal polarity axis;^17^ the basal surface is directly anchored on the basement membrane through integrins,^18^ and the apical surface is facing the lumen and is orientated away from the ECM.^19, 20^ Tight junctions^21^ provide a cell with different functionality between apical and basolateral membrane domains and help maintain apicobasal polarity as intramembranous diffusion of proteins and lipids is prohibited.^22, 23^ The basal/basolateral localization of polarity complex PAR-1b in the salivary gland further enforces the polarized organization.^24^ The correct polarization ensures the directional protein transport and deposition. For example, α-amylase is secreted to the apical surface, whereas laminins and type IV collagens are deposited to the basal side (Figure 1B).^23^

By regulating cytoskeleton dynamics, Rho family proteins mediate the positioning of the adherens junction and reshape the migrating cell, thereby contributing to the establishment and maintenance of epithelial polarization.^25^ As a Rho family effector, ROCK has been shown to control epithelial polarity. It is known that Y27632 inhibits RhoA downstream effectors ROCK-I and ROCK-II. Madin-Darby canine kidney (MDCK) cells expressing dominant-negative Rac1N17 or treated with the anti-β1-integrin antibody exhibited an inversion of polarity in three-dimensional (3D) collagen gel culture; ROCK inhibition led to the restoration of the correct polarity and formation of normal cysts.^26^ The mechanical state of the epithelial cells and their ECM can also influence cell polarity. A recent study shows that MDCK cells cultured on a soft substrate (1 kPa) were correctly polarized, but those on a stiffer one (>10 kPa) remained unpolarized.^27^ Interestingly, integrin inhibition allows for polarization on the supraphysiological substrates. Because ROCK inhibition is known to suppress myosin-mediated cell contractility and destabilize F-actin,^28, 29^ it is not surprising that ROCK inhibition led to a decrease in the stiffness of cancer cells grown in 3D collagen matrix.^30^ We speculate that ROCK-induced cytoskeletal tension in hS/PCs in 3D HA gels may contribute to the establishment of multicellular hS/PC spheroids that are inversely polarized. In addition to intrinsic cellular programs, the properties of the synthetic ECM, i.e., covalently crosslinked hydrogel network, can affect the stiffness of the resident cells. In fact, degradation-mediated cellular traction, not the particular shape of the cells, directs stem cell fate in covalently crosslinked 3D hydrogels.^31^ We reasoned that cell-mediated matrix degradation can cooperate with ROCK signaling to control hS/PC polarization in 3D HA gels.

Herein, we cultured hS/PCs in cell-adhesive HA gels with varying percentages of matrix metalloprotease (MMP)-cleavable crosslinks (nondegradable: 0DEG, 50% degradable: 50DEG, and 100% degradable: 100DEG, Figure 1A). We demonstrated that ROCK inhibition with Y27632 led to the establishment of acini-like multicellular structures containing a defined lumen lined with correctly polarized cells. The polarized structures showed apical localization of a Golgi marker, basal presentation of β1 integrin and basement membrane proteins, and apical and lateral localization of tight junction protein zonula occludens-1 (ZO-1). Noteworthy, Y27632-mediated polarization was matrix-dependent. Inhibition of ROCK in 0DEG and 50DEG cultures led to proper polarization, whereas inhibition of ROCK in 100DEG cultures led to cell scattering and loosening of the multicellular structures. Luminal secretion of α-amylase granules after neurotransmitter stimulation further confirmed the establishment of properly polarized and tightly sealed structures. Finally, we performed mitochondria-tracking microrheology analyses to examine how intracellular mechanics and cell stiffness dictate hS/PC polarization. This study demonstrates the importance of cell-mediated optimum matrix remodeling and cellular stiffness in salivary multicellular spheroid polarization.

## MATERIALS AND METHODS

### Synthesis of Hydrogel Precursors

Thiolated HA with ∼60% thiol incorporation was synthesized following our reported procedures.^8, 11, 32^ Maleimide functionalized peptide crosslinker that was susceptible [GIW-bisMI, GK(MI)RDGPQ<u>G↓IW</u>GQDRK(MI)G], or resistant [GIQ-bisMI, GK(MI)RD<u>GIQ</u>QWGGPDRK(MI)G] to MMP cleavage, as well as maleimide-functionalized cell adhesive peptide (RGD-MI, MI-GGG<u>RGD</u>SPG) were synthesized and purified as previously described.^11^ Peptide purity and mass were verified by UPLC and ESI-MS. All hydrogel precursors were stored as a lyophilized powder at −20 °C.

### 3D Culture of hS/PCs

Following our reported procedures,^8, 11, 12, 33^ hS/PCs were isolated from the parotid tissues of consented patients undergoing parotidectomy with human protocols approved by Christiana Care and Thomas Jefferson University. Cells at passages between 4-8 were used. hS/PCs were suspended in a 2 wt% HA-SH solution at pH 6.4. Next, RGD-MI and GIW-bisMI and/or GIQ-bisMI solutions were added. The final hS/PC concentration in all constructs was 3×10^6^ cells/mL. Varying the molar ratio of GIW-bisMI and GIQ-bisMI (0/1, 1/1, 1/0) while maintaining the overall concentration of peptide crosslinkers constant afforded constructs that were non-degradable (0DEG), 50% degradable (50DEG), and 100% degradable (100DEG). In all cases, the RGD concentration was maintained at 3 mM. HepatoSTIM (Corning) supplemented with epidermal growth factor (EGF), pen-strep, and amphotericin B, with or without the ROCK inhibitor Y27632 (20 μM, Abcam) was added 5 min after gel fabrication. The cellular constructs were cultured for 15 days with media refreshment every three days. The constructs were inspected using a phase contrast light microscope (Nikon Eclipse Ti series) for the development of multicellular structures.

### Immunocytochemistry

The cellular constructs were fixed in 4% paraformaldehyde (PFA, Sigma Aldrich) for 30 min and permeabilized in 0.2% Triton (Sigma Aldrich) for GM130 and α-amylase or 0.05% saponin (Sigma Aldrich) for ZO-1 and integrin β1, both with 3% (w/v) bovine serum albumin (BSA, Cell Signaling Technology), for 16 h at room temperature. No permeabilization was performed for laminin α1, collagen IV, and fibronectin. Constructs were incubated in primary antibody solutions for 24 h at room temperature. After two PBST (1× PBS with 0.05% Tween 20 Sigma Aldrich) washes (15 min each), samples were incubated with PBS for 16 h. Constructs were then incubated with the secondary antibody (1:200) along with Alexa Fluor™ 568 phalloidin (Life Technologies, 1:250) in the respective permeabilizing/blocking solution for 24 h at room temperature. The cell-laden constructs were washed thrice with PBST for 20 min each. The constructs were then stained with DAPI (Life Technologies, 1:1000) in PBST for 30 min at room temperature and washed with PBST thrice for 20 min each. To characterize amylase secretion, 50 µM isoproterenol was added to the media, and the construct was maintained at 37 °C with 5% CO_2_ for 1 h before PFA fixation and subsequent staining for α-amylase, F-actin, and nuclei. See Table S1 for antibody information and dilution conditions. Confocal microscopy imaging was performed using a Zeiss LSM 880 equipped with an Airyscan detector in Fast Airyscan mode. Unless specified, images were presented as single slices. Maximum intensity projections were obtained as z-stacks of 50 μm using ImageJ/Fiji.

### Image Analysis

Using the ZEN Black software, each slice in the z-stack image was manually inspected for the presence of a cell-free hollow center, a hollow center with apically localized GM130, and a hollow center with α-amylase stains. Normalization of the number of spheroids with a hollow center to the total number of spheroids yielded percent spheroids with lumens. Normalization of the number of lumen-containing spheroids with apically localized GM130 to the total number of spheroids with lumens yielded the percentage of spheroids that were properly polarized. A total of 30 images (40×) per condition were analyzed. All spheroids in each image were analyzed. Supplementary videos 1-6 show confocal z-stack movie of hS/PC spheroids stained with F-actin (red) and nuclei (blue).

### Quantitative Polymerase Chain Reaction (qPCR)

The cell-laden hydrogel constructs were snap-frozen in isopropanol and dry ice and crushed with a pestle. One milliliter Trizol solution (Invitrogen) was added to the gel slurry and mixed with the pipette. This solution was incubated in an Invitrogen™ Phasemaker™ Tube (Thermo Fisher Scientific) for 3 min at room temperature. After the addition of 200 μL chloroform, the tube was inverted by hand and rotated by centrifugation at 12,000 ×g at 4°C for 15 min. The aqueous layer was collected in a separate Eppendorf tube, and its volume was noted. After adding an equal amount of 100% ethanol, the solution was transferred to RNA extraction columns (Zymo Research), and the solvents were removed by centrifugation at 16,000 ×g for 30 s at room temperature. After adding 400 μL RNA prep buffer (Zymo Research), the column was centrifuged at 16,000 ×g for 30 s. After adding 700 μL of wash solution (Zymo Research), the column was centrifuged again at 16,000 ×g for 30 s to remove the solution. This step was repeated one more time. Upon addition of the wash buffer (400 μL), the column was centrifuged at 16,000 ×g for 2 min. Immediately after the addition of 6 μL DEPC water (Fisher Scientific), the column was centrifuged at 16,000 ×g for 2 min, and aqueous eluents were collected. This protocol yielded high-purity mRNA with absorbance ratios at 260/280 nm > 2.01 and 260/230 > 1.95, as analyzed by a NanoDrop 2000 Spectrophotometer (Nanodrop Technologies).

The mRNA was reverse transcribed to cDNA using the QuantiTect® Reverse Transcription Kit (QIAGEN) following the manufacturer’s protocol. Using an Applied Biosystems 7300 real-time PCR machine, sequence-specific amplification and detection were performed. Thermal cycling was performed with 1 cycle at 95 °C for 10 min, followed by 40 cycles at 95 °C for 15 s each, and 1 cycle at 60 °C for 1 min. PCR reactions were prepared by combining Power SYBR™ green PCR master mix (Applied Biosystems), cDNA, and target-specific primers for a total volume of 20 μL per reaction. Primers were ordered from Integrated DNA Technologies, and the complete primer sequences are available in Table S2. Glyceraldehyde 3-phosphate dehydrogenase *(GAPDH)* was used as the housekeeping gene. Cycle threshold values were generated using 7300 System SDS RQ Study software version 1.4 (Applied Biosystems). The obtained C_T_ values were normalized to *GAPDH*. The fold changes were calculated using the ΔΔC_T_ method. Three biological replicates are reported from three technical replicates measured in duplicate.

### Mitochondria-Tracking Microrheology

On days 1 and 15, the media was supplemented with 500 nM of MitoTracker green (Life Technologies) and the culture was maintained at 37°C and 5% CO_2_ for 1 h. Time-lapse images of stained mitochondria were acquired using a Zeiss LSM 880 confocal microscope with an Airyscan detector and a 40× oil immersion objective. During live cell imaging, constructs were maintained at 37°C with 5% CO_2_. ZEN Black software was used to capture the time-lapse images. Cells were located and tracked for 105 s. At least 60 time-lapse images were captured for each condition at each time point. Images were acquired from three independent biological repeats. Using TrackMate, a Fiji Image J plugin, mitochondrial fluctuations were extrapolated from the time-lapse images as particle trajectories. The particle trajectories were converted to time-dependent mean square displacement (MSD) using MATLAB (Math Works, Inc., 2020) and MSD-Analyzer according to:

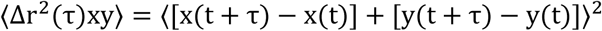

where ⟨Δr^2^(τ)xy⟩ is the ensemble-averaged MSD, τ is the delay time between the first and last image frame analyzed, and x(t) and y(t) are the spatial coordinates of a particle position at the time t. For viscoelastic materials, the MSD scales nonlinearly with the delay time τ according to a power-law relationship, i.e. ⟨Δr^2^(τ)⟩ ∼ τα. The power-law coefficient α = ∂ln 〈Δr^2^ (τ)〉/ ∂ln (τ) represents the slope of the logarithmic MSD-τ curve.

### Statistical Analysis

Statistical analysis was conducted using JMP Pro 15 (SAS Institute Inc.). Error bars represent the standard error of the mean (SEM). Comparisons were made based on student’s t-test or one-way ANOVA with Tukey’s HSD post hoc. A *p*-value of less than 0.05 was considered statistically significant.

## RESULTS

### ROCK Inhibition Promotes Cell Polarization in Hydrogels with Optimal Degradability

HA-based, cell adhesive hydrogels with a similar initial storage modulus (∼200 Pa) but varying susceptibility to cell-mediated proteolytic degradation (non-degradable-0DEG, 50% degradable-50DEG, and 100% degradable-100DEG) were used as the synthetic ECM for the 3D culture of hS/PCs with or without the ROCK inhibitor Y27632 (Figure 1A). hS/PCs encapsulated as single cells grew into multicellular structures under all conditions. Without Y27632, spheroid circularity and compactness decreased as matrix degradability increased.^11^ Supplementation of the culture media with Y27632 led to the formation of multicellular spheroids with hollow lumens in 0DEG and 50DEG hydrogels (Figure 2A). On average, 64% and 80% of spheroids developed in 0DEG and 50DEG matrices contained a hollow lumen (Figure 2B). However, multicellular structures detected in 100DEG gels were irregularly shaped and scattered. Only 8% of structures in 100DEG gels were lumenized, significantly (*p* < 0.01) lower than the 0DEG and 50DEG constructs. Without ROCK inhibition, spheroids with a central lumen were rarely seen under 0DEG and 50DEG conditions.

**Figure 2.**
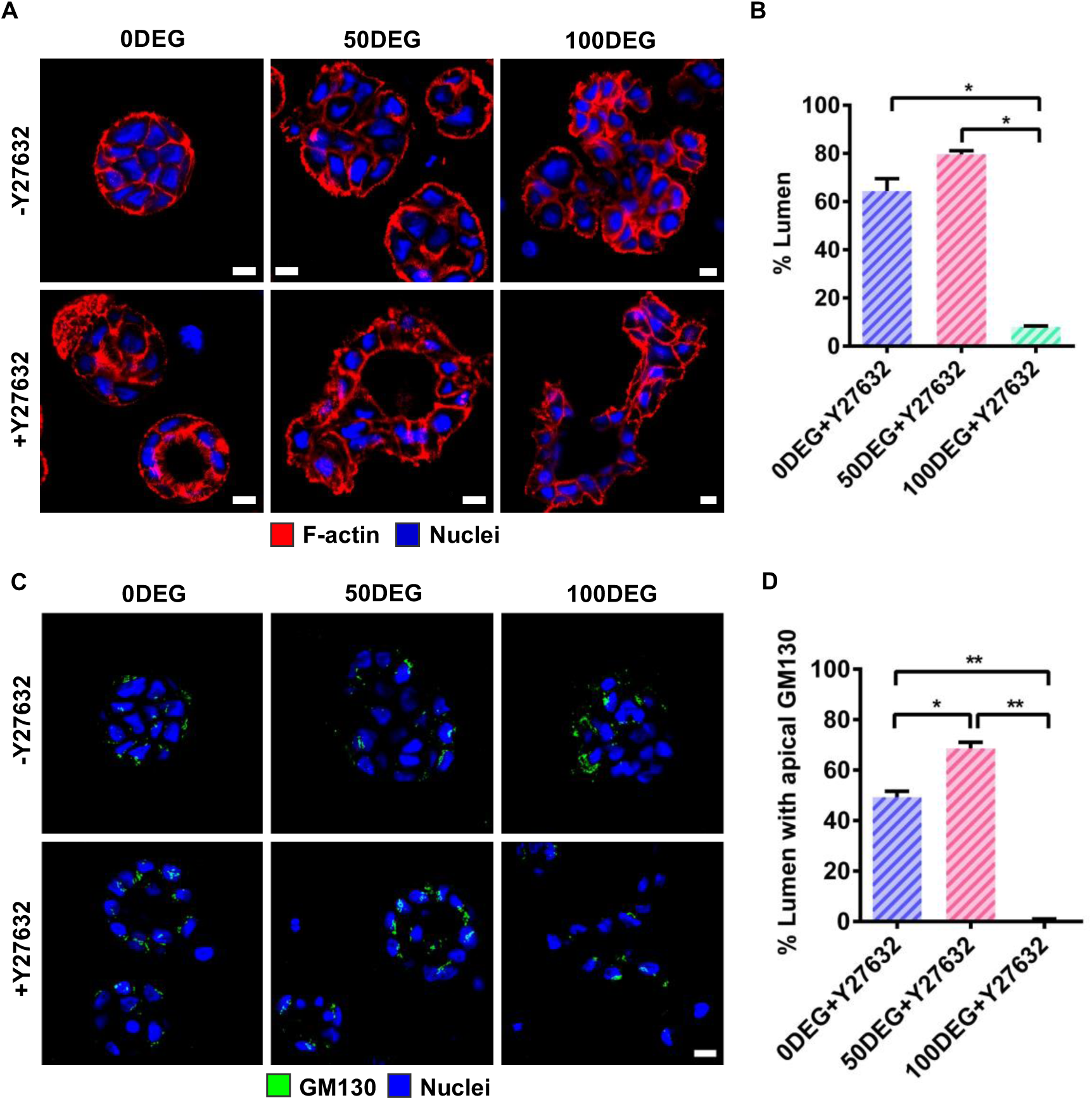
ROCK inhibition in 0DEG and 50DEG cultures leads to the establishment of lumen-containing spheroids with apicobasal polarization. **(A)** Confocal images of day 15 cultures stained for F-actin (red) and nuclei (blue). Scale bar: 10 µm. **(B)** Quantification of spheroids with lumens in Y27632-treated cultures on day 15. Error bars represent SEM. One-way ANOVA post hoc Tukey; * indicate *p* < 0.05. **(C)** Immunofluorescent images depicting GM130 (green) localization. Cell nuclei were stained blue with DAPI. Apical localization of GM130 suggests correct polarization. Scale bar: 10 µm. **(D)** Quantification of lumens with apical staining of GM130. Day 15 constructs were analyzed. Error bars represent SEM. One-way ANOVA post hoc Tukey; * and ** indicate *p* < 0.05 and 0.01, respectively.

It is known that the Golgi apparatus localizes to the apical side of the nucleus in a manner depending on the polarized organization of the microtubule network (Figure 1B).^34^ In the absence of Y27632, the GM130 signal was randomly localized throughout the spheroid in all three types of cellular constructs. However, GM130 was detected between the nucleus and the apical membrane in Y27632-conditioned 0DEG and 50DEG cultures, indicating that the structures were correctly polarized (Figure 2C). Quantitatively (Figure 2D), 50DEG cultures contained approximately 69% correctly polarized spheroids, significantly (*p* < 0.05) higher than those found in the 0DEG cultures (49%). This contrasts sharply with the 100DEG gels where less than 1% of structures contained correctly polarized lumens.

Located in the basolateral domain in the salivary gland, PAR-1b promotes laminin accumulation, mediates microtubule orientation, and controls the asymmetric distribution of cell surface proteins.^35^ In the engineered constructs, however, PAR-1b was detected around individual cells, colocalized with cortical F-actin. The staining patterns were similar between the Y27632-free and Y27632-treated samples, although the latter lacked a lumen (Figure 3A). In normal epithelial tissues, tight junction proteins connect the cells to seal the structure and act as a boundary between the apical and basolateral membrane domains to induce cell polarization; cells are anchored through integrins to the basement membrane at the basal surface (Figure 1B). These cell-matrix and cell-cell interactions provide directional cues to establish polarity and promote cell survival, proliferation, and differentiation.^36^ ZO-1 (or *TJP1*) is an intracellular scaffolding protein that plays a vital role in tight junction protein assembly. Y27632-free cultures displayed basal and lateral staining of ZO-1 (Figure 3B). Supplementation of the 50DEG culture with the ROCK inhibitor led to apical and lateral localization of ZO-1. The spatial redistribution of ZO-1 at the protein level caused by Y-27632 was accompanied by a significant reduction (1.4-fold, *p* < 0.01) in the expression of *TJP1* at the transcript level for 50DEG cultures (Figure S1A). For 0DEG and 100DEG controls, however, Y27632 treatment did not alter *TJP1* expression significantly. A similar gene expression profile was detected for occludin (*OCLN*), another tight junction protein (Figure S1B). ROCK inhibition resulted in a 2.6-fold (*p* < 0.01) and a 2.1-fold (*p* < 0.05) decrease in *OCLN* expression for 50DEG and 0DEG cultures, respectively, but no changes were detected for the 100DEG counterparts.

**Figure 3.**
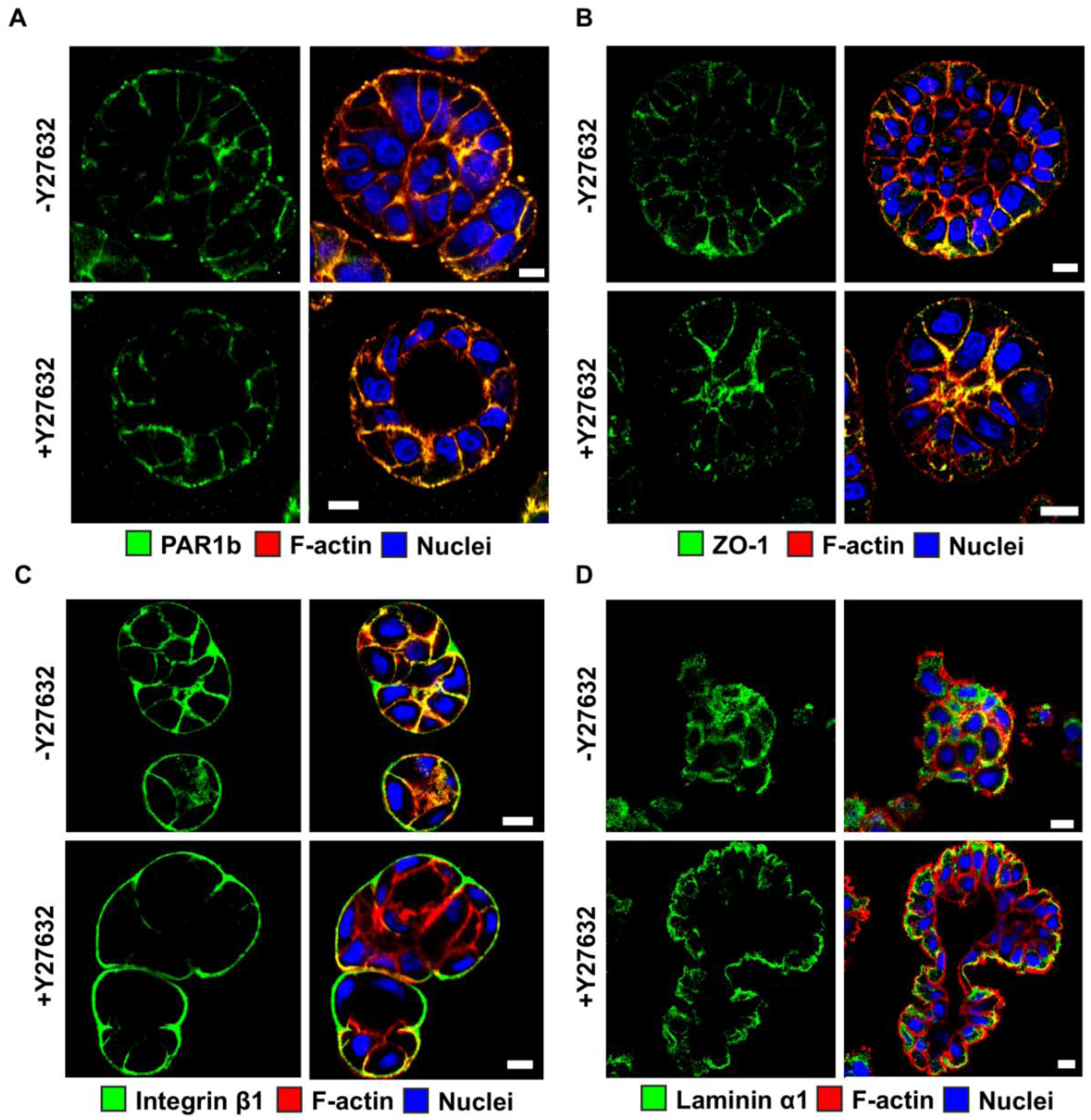
ROCK inhibition in 50DEG cultures leads to spatial localization of membrane proteins (A-C) and a basement membrane protein laminin α1 (D). **(A)** Immunofluorescent images of day 15 50DEG cultures stained for PAR-1b (green). F-actin and nuclei are stained red and blue, respectively. Scale bar: 10 µm. **(B)** Immunofluorescent images of day 15 50DEG cultures stained for ZO-1 (green). F-actin and nuclei are shown in red and blue, respectively. Scale bar: 10 µm. **(C)** Immunofluorescent images of day 15 50DEG cultures stained for integrin-β1 (green). F-actin and nuclei are shown in red and blue, respectively. Scale bar: 10 µm. **(D)** Immunofluorescent images of day 15 50DEG cultures stained for laminin α1 (green). F-actin and nuclei are shown in red and blue, respectively.

Immunocytochemical analysis revealed a significant difference in the spatial localization of integrin β1 (Figure 3C). Without ROCK inhibition, integrin β1 was found cortically around individual cells overlapping with F-actin. In the presence of Y27632, however, integrin β1 was restricted to the basal position around the entire spheroid and did not overlap with the F-actin signal. A similar differential integrin β1 localization was observed in 0DEG cultures (Figure S2A). However, integrin β1 was sequestered at the periphery of individual cells in 100DEG cultures independent of Y27632 treatment. At the mRNA level, Y27632 treatment resulted in a significant decrease (2.3-fold, *p*< 0.01) in *ITGB1* expression of in 50DEG cultures, but the expression level was comparable for other cultures with or without Y27632 (Figure S2B).

Basement membrane proteins secreted by epithelial cells at their basal side define the basal polarity.^20, 37^ The staining patterns for 50DEG cultures for laminin α1 differed considerably under the prescribed 3D culture conditions (Figure 3D). Laminin α1 was detected throughout the spheroids at individual cell peripheries for Y27632-free cultures. However, robust continuous staining delineating the border of the entire multicellular structure was observed in cultures supplemented with Y27632. A similar differential laminin α1 localization was observed in 0DEG cultures (Figure S3A). Again, laminin α1 was found intercellularly in 100DEG cultures irrespective of Y27632 treatment. Y27632 treatment did not result in a significant change in the expression of *LAMA1* for 0DEG and 50DEG cultures but did give rise to a significant increase (2.5-fold, *p* < 0.05) in 100DEG cultures (Figure S3B).

Immunostaining for collagen IV (Figure S4) revealed a similar trend to laminin α1, with the protein deposited at the basal side of multicellular structures in 0DEG and 50DEG cultures with Y27632. Without ROCK inhibition, collagen IV signals were predominately intracellular, irrespective of the hydrogel composition. ROCK inhibition stimulated the deposition of collagen IV in the extracellular space for all three types of cultures, albeit at an ectopic location for 100DEG cultures. As a putative cleft initiator, the ECM protein fibronectin directs the inward translocation of cells to promote branching during development.^38^ In the absence of Y27632, fibronectin was expressed intracellularly (Figure S5A). ROCK inhibition led to the extracellular deposition of fibronectin with varying spatial localization. At the transcript level, ROCK inhibition resulted in a significant decrease in FN1 expression in 0DEG (4-fold, *p* < 0.05) and 100DEG (2.6-fold, *p* < 0.05) cultures (Figure S5B). Thus, the expression levels of genes encoding these proteins do not correlate positively with their spatial localization at the protein level.

### ROCK Inhibition Stimulates the Development of Pro-Acinar Progenitor Cells in 50DEG

Day 15 50DEG constructs were evaluated for the expression of stem/progenitor markers at the transcript level by RT-qPCR (Figure 4A). Keratins 5 and 14 are dimerizing partners co-expressed by stem cells in developing salivary glands.^39^ Y27632 treatment resulted in a 9.4-fold (*p* < 0.005) and 2.1-fold (*p* < 0.005) increase in *KRT5* and *KRT14* expression, respectively. Compared to the control culture, the Y27632-conditioned culture exhibited a 2.1-fold (*p* < 0.05) higher expression of *MYC,* a transcription factor required to maintain the stem/progenitor pool.^40^ Receptor tyrosine kinase KIT and its ligand KITLG, also known as stem cell factor (SCF), play a critical role in the growth and survival of salivary gland epithelial cells.^40, 41^ ROCK inhibition upregulated *KITLG* expression (2.2-fold, *p* < 0.01), with a concurrent downregulation of *KIT* expression (16.4-fold, *p* < 0.001). Immunostaining for K14 revealed abundant filamentous structures inside all constituent cells in individual multicellular structures in Y27632 cultures. However, K14 signals were more pronounced in cells located at the outer surface of the spheroids in Y27632-free cultures (Figure 4B).

**Figure 4.**
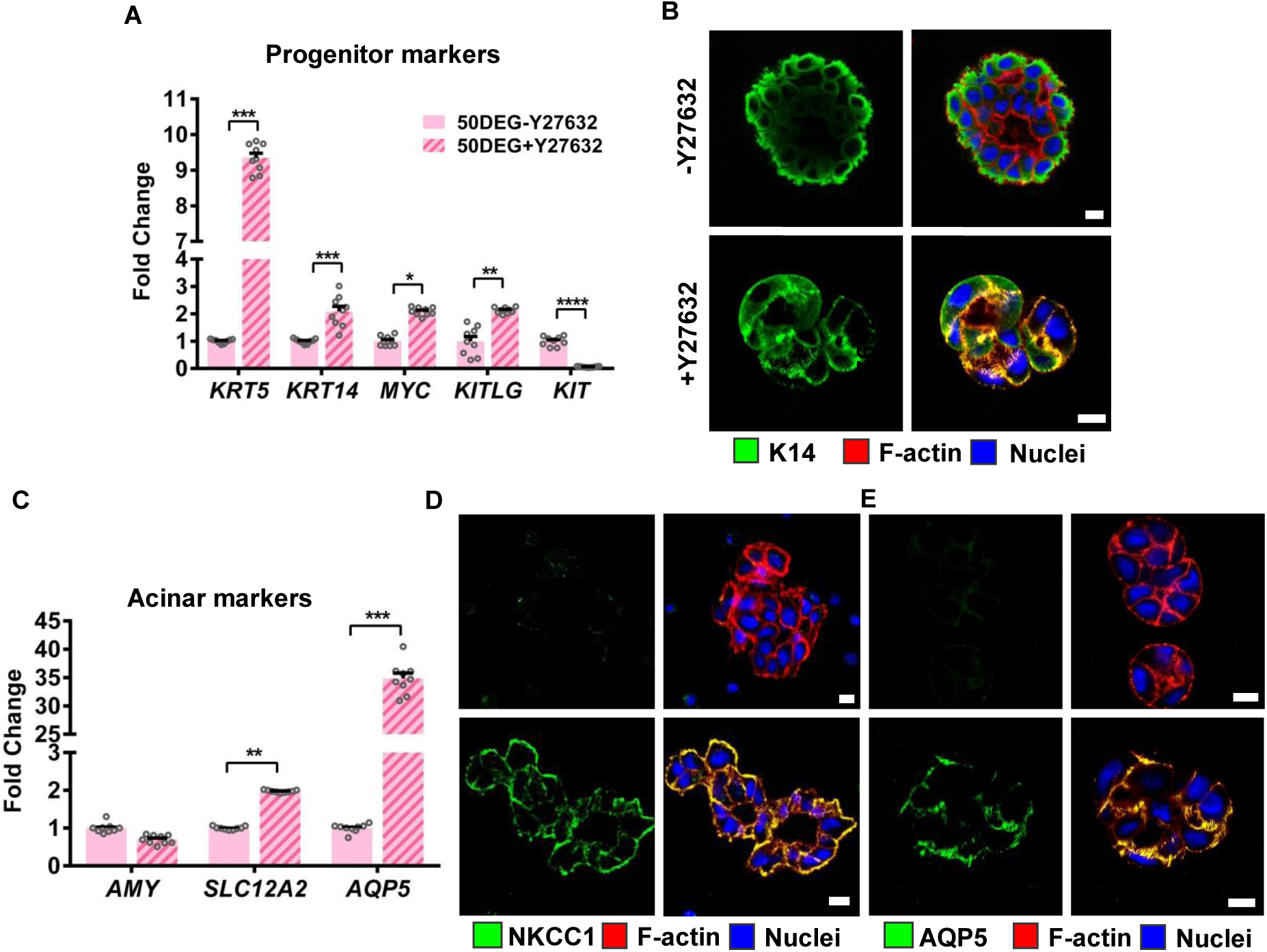
ROCK inhibition in 50DEG cultures increases the expression of stem/progenitor and acinar markers. **(A)** RT-qPCR analyses of day 15 50DEG constructs cultivated with or without Y27632 for salivary gland stem (*MYC*, *KITLG*, *KIT*) and progenitor (*KRT5*, *KRT14*) cell markers. Error bars represent SEM. Student’s t-test was performed, and *, **, ***, and **** indicate *p* < 0.05, 0.01, 0.005, and 0.001, respectively. **(B)** Confocal images of day 15 50DEG constructs stained for progenitor marker K14 (green), F-actin (red), and nuclei (blue). Scale bar: 10 µm. **(C)** RT-qPCR analyses of day 15 50DEG constructs cultivated with or without Y27632 for salivary gland acinar cell markers (*AMY, SLC12A2, AQP5*). Error bars represent SEM. Student’s t-test was performed, and *, **, and *** indicate p < 0.05, 0.01, and 0.005, respectively. **(D)** Confocal images of day 15 50DEG constructs stained for acinar maker NKCC1 (green), F-actin (red), and nuclei (blue). Scale bar: 10 µm. **(E)** Confocal images of day 15 50DEG constructs stained for acinar makers AQP5 (green), F-actin (red), and nuclei (blue). Scale bar: 10 µm

Y27632 also enhanced *KRT5* and *KRT14* expression in 0DEG cultures although the actual fold changes were lower as compared to the 50DEG cultures (Figure S6). In 100DEG cultures, ROCK inhibition led to significant down-regulation of *MYC* (2.1-fold, *p* < 0.01) and *KIT* (17.4-fold, *p* < 0.005) expression, but had no effects on *KRT5, KRT14 and KITLG.* Interestingly, in the non-degradable hydrogels, Y27632 treatment did not significantly change the cellular expression of *KIT*. However, when the hydrogel became proteolytically degradable, ROCK inhibition substantially decreased *KIT* expression. Like the 50DEG conditions, 0DEG and 100DEG cultures, irrespective of Y27632 treatment, were stained positive for K14 (Figure S7). Without Y27632, K14 staining was stronger in cells located at the border of the spheroids in 100DEG constructs. In 0DEG cultures without Y27632, cells in the spheroid core were also K14-positive. Taken together, Y27632 was most effective in maintaining stem/progenitor phenotype when cells were cultured in 50DEG gels.

Next, we examined the expression of acinar markers *AMY*, *SLC12A2,* and *AQP5* at the mRNA level (Figure 4C) by 50DEG cultures. Y27632 treatment resulted in a small, non-significant decrease in *AMY* expression. Expression of *SLC12A2* was increased by 2-fold, significantly (*p* < 0.01) higher than the untreated controls. ROCK inhibition yielded 34.8-fold increase (*p* < 0.005) in the expression of *AQP5*, a transmembrane channel protein facilitating water transport across the cell membrane and a target for gene therapy for the treatment of xerostomia. A comparison with 0DEG and 100DEG cultures again revealed that the regulatory effects of Y27632 manifested most profoundly when hS/PCs were cultured in the 50DEG matrix (Figure S8).

NKCC1 is a Na-K-Cl cotransporter that aids in the active transport of these ions into cells and is indispensable for maintaining secretory functions in salivary gland acinar cells.^42^ Cultures maintained in 50DEG without ROCK inhibition were stained negative for NKCC1, whereas those maintained with Y27632 were stained brightly for NKCC1 (Figure 4D). Similarly, 50DEG/Y27632 cultures were stained robustly for AQP5, whereas only faint background signals were detected for 50DEG cultures without Y27632 (Figure 4E). For 0DEG constructs, Y27632 treated samples exhibited enhanced staining patterns for NKCC1 and AQP5 compared to the untreated controls. For 100DEG constructs, however, the staining pattern and intensity were similar between cultures with or without Y27632 treatment (Figures S9, S10).

Characterization of ductal markers at the transcript level (Figure S11) revealed that supplementation of the 50DEG cultures with Y27632 led to a significant decrease in the expression of *TFC2PL1* (2.4-fold, *p* < 0.05) and *KRT7* (2-fold, *p* < 0.01). ROCK inhibition did not significantly alter the expression of *MUC1.* For 0DEG cultures, ROCK inhibition did not significantly alter the expression of *TFCP2L1*, *KRT7*, or *MUC1*. For 100DEG cultures, Y27632 treatment resulted in a significant decrease in *TFCP2L1* (3.7-fold, *p* < 0.01), a significant increase in *KRT7* (1.9-fold, *p* < 0.05), and a significant reduction in *MUC1* expression (2.3-fold, *p* < 0.05). Overall, ROCK inhibition in 50DEG cultures promoted acinar differentiation and suppressed differentiation towards the ductal lineage.

### ROCK Inhibition Mediates *ROCK Expression* and Enables Vectorial Saliva Secretion in 50DEG cultures

The expression levels of *ROCK1* and *ROCK2* under different culture conditions were analyzed by RT-qPCR. In the absence of Y27632, increasing matrix degradability resulted in a significant increase in the expression of both *ROCK1 and ROCK2* (Figure 5A); *ROCK1* expression level was the highest in 100DEG cultures, whereas *ROCK2* expression level was comparable in 50DEG and 100DEG. ROCK inhibition led to a significant (p < 0.05) decrease (1.3-fold) in the expression of *ROCK1* only in the 50DEG matrix. ROCK inhibition did not significantly alter the expression of ROCK2 across different gel formulations (Figure 5B). Serous acinar cells are abundant in the parotid gland and have well-developed cytoplasmic organelles, including rough endoplasmic reticulum, Golgi apparatus, and secretory granules. The granules contain proteins such as α-amylase that are secreted in the central lumen of the acinus.^11^ To assess the secretory function of multicellular spheroids developed in 50DEG/Y27632 cultures, isoproterenol, a β-adrenergic agonist, was used to stimulate protein exocytosis. If the structures are correctly polarized, exocytosis should occur towards the apical side of the lumen (Figure 1B). Under control conditions without Y27632, α-amylase was secreted outside the spheroids towards the basal side (Figure 5C), indicating inverse polarization. Contrarily, amylase granules were secreted into the lumen from the apical side of the spheroids in Y27632-conditioned cultures, confirming correct apicobasal polarity. Qualitatively, the amount of α-amylase detected in Y-27632-conditioned cultures appeared higher too.

**Figure 5.**
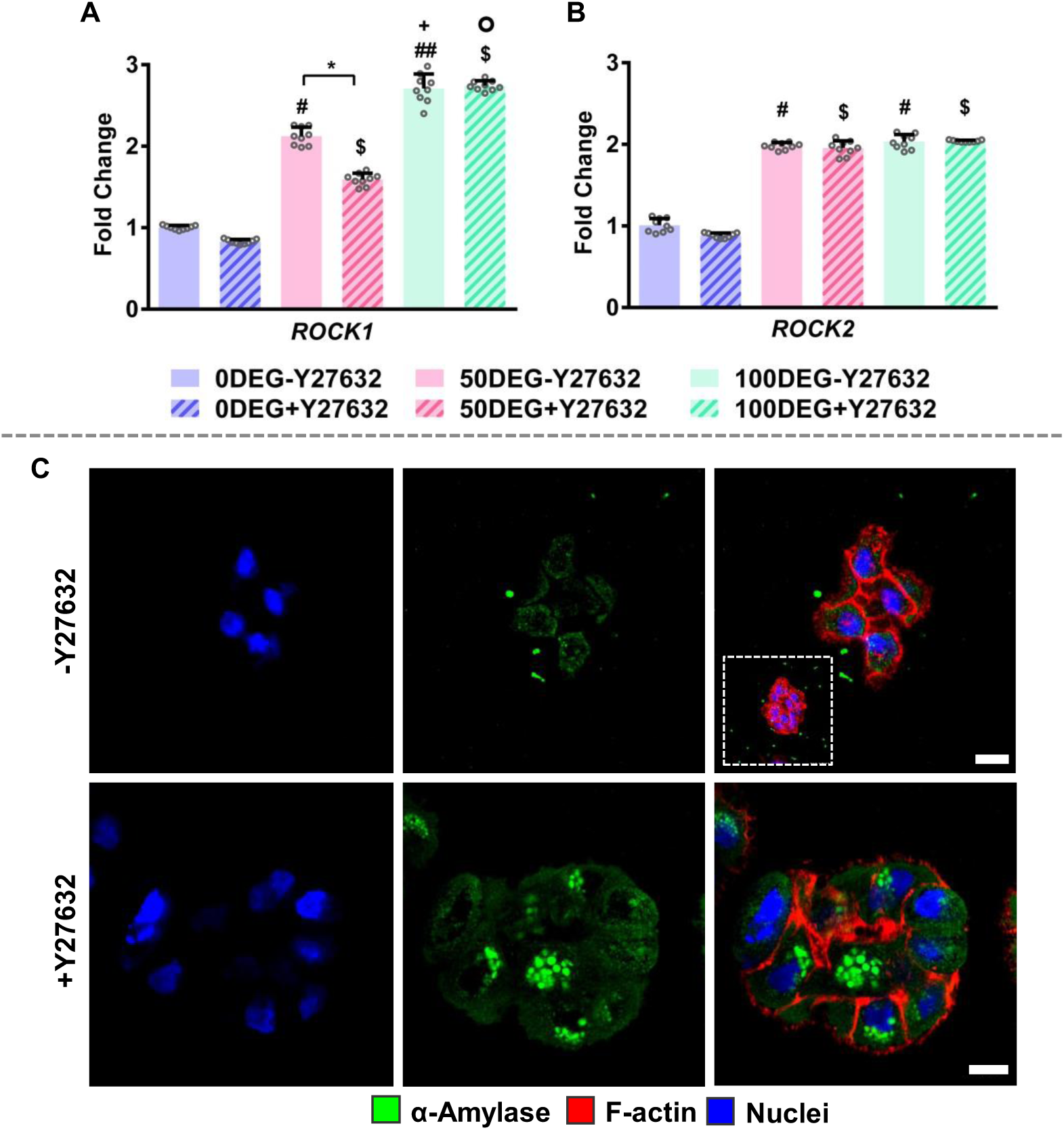
ROCK inhibition in 50DEG suppresses *ROCK1* expression and stimulates vectorial secretion of α-amylase in 50DEG cultures. **(A)** RT-qPCR analyses of day 15 constructs maintained with or without Y27632 for the expression of *ROCK1*. **(B)** RT-qPCR analyses of day 15 constructs maintained with or without Y27632 for the expression of *ROCK2*. Expression was normalized to Y27632-free 0DEG constructs. Error bars represent SEM. One-way ANOVA with post hoc Tukey’s test was performed; * indicates *p* < 0.05 between 50DEG cultures with and without Y27632 treatment. ^#^ indicates *p* < 0.05 for Y27632-free 0DEG and 50DEG cultures. ^##^ indicates *p* < 0.01 for Y27632-free 0DEG and 100DEG cultures. *^+^* indicates *p* < 0.01 for Y27632-free 50DEG and 100DEG cultures. ^$^ indicates *p* < 0.05 for Y27632-treated 0DEG and 50DEG cultures. ° indicates *p* < 0.01 for Y27632-conditioned 50DEG and 100DEG. **(C)** Confocal images (single slices, with the insert showing a maximum intensity projection) of 50DEG constructs stained for amylase (green), F-actin (red), and nuclei (blue) after 15 days of culture with or without Y27632, followed by 1-h stimulation with isoproterenol. Scale bar: 10 µm.

### Matrix Degradability and ROCK Inhibition Influence hS/PC Stiffness

Particle tracking microrheology was employed to examine cellular mechanics under different culture conditions. Instead of exogenous particles ballistically injected into cells, here mitochondria were used as endogenous tracer particles to monitor cellular mechanics non-invasively (Figure S12). A lower ensemble-average MSD represents constrained mitochondrion fluctuations and stiffer solid material, whereas a high MSD indicates greater particle motility and a more fluid-like material. The power-law coefficient α helps classify the motion of the particles: α ≈ 1 corresponds to diffusive motion, such as thermal fluctuations in Newtonian fluids, and α ≈ 0 represents constrained, sub-diffusive motion, such as thermal fluctuations in an elastic material. The mitochondrial MSD represents the viscoelastic intracellular properties for short delay times, whereas active intracellular motor-driven processes dominate the MSD for long delay times. For a non-equilibrium system such as the intracellular environment, molecular motor activity results in additional internal fluctuations beyond thermal fluctuations, leading to increased MSD and α at relatively longer time scales. Here, data acquired at short delay times (0–1 s) were used to extract cell stiffness.^30^

Shown in Figures 6A and 6C are MSD curves (i-iii) and the average MSD values at a delay time of 50 ms for 0DEG, 50DEG, and 100DEG cultures with or without Y27632 one day after cell encapsulation when hS/PCs remained as single cells under all culture conditions. In all three types of gels, adding Y27632 led to a significant (*p* < 0.01 for 0DEG and 100DEG; *p* < 0.05 for 50DEG) increase in the MSD values. Without Y27632, transitioning from non-degradable gels to 100% degradable gels significantly (*p* < 0.01) reduced the MSD values. Thus, increasing matrix degradability rendered cells stiffer, and inhibiting ROCK made cells softer. However, on day 15, when hydrogel-derived multicellular structures varied significantly in size and morphology across cultures, the average MSD values measured for all cultures collapsed to a similar value, irrespective of the matrix degradability and whether Y27632 was present (Figure 6B, 6D, *p* > 0.05). We also examined the power-law dependence of MSD at a time interval of 1 s (Table S3). On day 1, without Y27632, the α value was consistent across the three types of cultures, averaging 0.4-0.5. Similarly, the alpha values for Y27632-treated cultures were independent of matrix degradability, averaging around 0.3, consistently lower than those without ROCK inhibition. However, by day 15, the α values became similar. The decrease in α indicates that cells exhibited less fluid-like intracellular motions when treated with Y27632. Previous work on metastatic breast cancer cell lines encapsulated in collagen gels showed an increase in MSD accompanied by a decrease in α value.^30^

**Figure 6.**
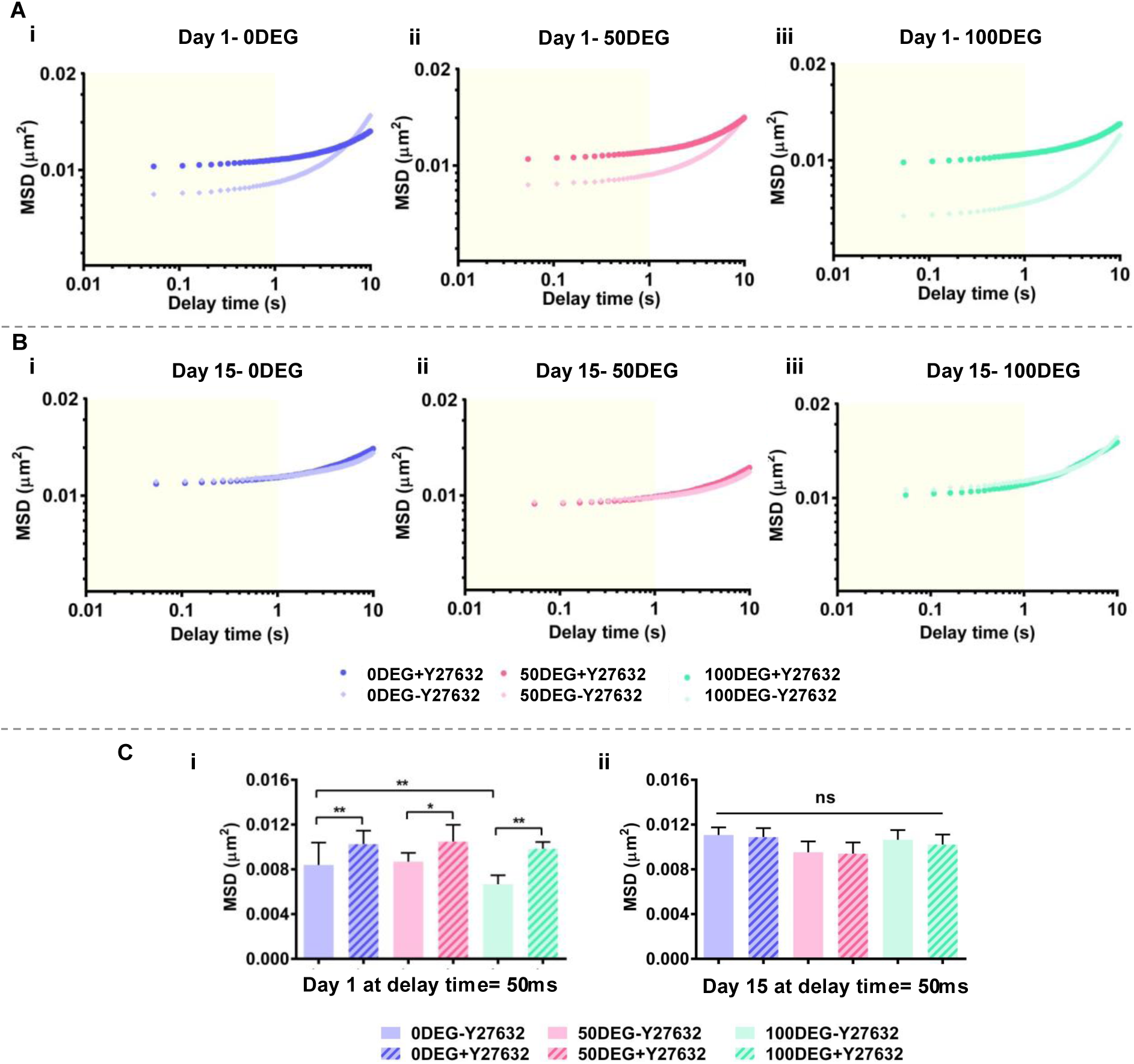
Matrix degradability and ROCK inhibition affect cell stiffness. **(A)** MSD curves of mitochondria in hS/PCs cultured in HA gels with varying degrees of degradability for 1 day with or without Y27632. **(B)** MSD curves of mitochondria particles in hS/PCs cultured in HA gels with varying degrees of degradability for 15 days with or without Y27632. **(C)** MSD values at a delay time of 50 ms on day 1 (i) and day 15 (ii) for hS/PCs cultured in HA gels with varying degradability with and without Y27632. Error bars represent SEM. One way ANOVA tests were performed; * and ** indicate *p* < 0.05 and 0.01, respectively. MSD data at a delay time of less than 1 ms (yellow shaded areas in A and B) represent passive particle diffusion. MSD data at longer delay times were not used due to the contribution of active intracellular motor-driven processes.

## DISCUSSION

A significant challenge in developing a tissue-engineered salivary gland using primary adult human salivary gland epithelial cells is establishing correctly polarized, acini-like multicellular structures for vectorial secretion.^14^ Although hS/PCs encapsulated as single cells in RGD-containing HA gels with increasing susceptibility to MMP-mediated degradation can form multicellular spheroids, in the absence of Y27632, apicobasal polarization was not observed. Culturing hS/PCs in 0DEG or 50DEG in the presence of Y27632 gave rise to multicellular structures that were correctly polarized. Apical-basal polarity was confirmed by the localization of Golgi marker GM130 between the nucleus and the apical surface, basal localization of basement membrane proteins, basal localization of integrin β1, apical-lateral localization of tight junction proteins, and secretion of α-amylase into the lumen. Notably, the localization patterns observed in our engineered gland are in good agreement with what is observed in healthy, adult human salivary glands.^43, 44^ On the one hand, a significant fraction (>60%) of multicellular structures developed in Y27632-conditioned 0DEG and 50DEG cultures are correctly polarized. On the other hand, ROCK inhibition in 100DEG led to considerable cell dissemination. As the matrix became progressively degradable, the multicellular structures became larger, less spherical, and more spread out, with or without Y27632.

ROCK1 inhibition causes activation of Rac1 to promote basal localization of integrins.^45, 46^ Basal localization of integrin β1 not only assists the organization of the basolateral surface^47, 48^ but also directs basal deposition of basement membrane proteins. Establishing a cohesive basement membrane surrounding the expanding spheroids is an essential indicator for correct polarization.^20^ Therefore, it is not surprising that β1 integrin ablation resulted in a loss of polarity, leading to defective arterial lumen formation^49^ and inverted polarity in glandular epithelium.^47^ Here, intracellular localization of basement membrane proteins (laminin α1 and collagen IV) in multicellular structures developed in all three types of gels without Y27632 and in Y27632-conditioned 100DEG cultures coincided with inverse polarization.

Previously, we showed that hS/PCs residing in cell-adhesive, MMP-degradable HA gels actively remodeled the matrix; as the matrix became progressively more degradable, protease secretion increased accordingly.^11^ Here, we show that matrix degradability correlates positively with the mRNA levels for *ROCK1* and *ROCK2*. Additionally, ROCK inhibition in 50DEG led to a significant decrease in *ROCK1* expression in 50DEG. Others show that interaction of integrins with ECM leads to ROCK activation, ROCK activation leads to ECM degradation,^50^ and inhibition of ROCK decreases the expression of MMPs.^51^ Thus, Y27632-treated to 50DEG cultures exhibit an optimal protease activity to promote the development of polarized structures. Here, 50DEG/Y27632 cultures exhibit a robust and cohesive basement membrane that is critically important for proper polarization.

Previously, we showed hS/PCs maintained in MMP degradable gels expressed a significantly higher level of KIT (an important stem/progenitor maker) than those grown in mechanically matched non-degradable gels. Upon ROCK inhibition, the downregulation of KIT was accompanied by the upregulation of key acinar markers (*AQP5* and *SLC12A2* by qPCR; AQP5 and NKCC1 by immunofluorescence), suggesting the differentiation of KIT+ cells to acinar cells; however, differentiation to the ductal cells was suppressed. Again, enhanced acinar differentiation was not observed in Y27632-conditioned 100DEG cultures. Our findings highlight the importance of optimal matrix degradability and ROCK inhibition to achieve lumen formation, apicobasal polarization, and acinar differentiation.

Our microrheology experiments revealed significant differences in the passive diffusion of mitochondrial particles in cells cultured under different conditions on day 1 when cells existed as single cells with intimate contact with the synthetic ECM. The MDS curves collapsed to a similar value by day 15 because the multicellular structures were heterogeneous and contained variable amounts of cell-cell and cell-ECM interactions, potentially negating the environmental effects since cells on the surface of the spheroids behave differently from those in the interior. We emphasize that the cell fate (polarity and phenotype) was determined early on when hS/PCs were dispersed in the matrix as single cells.

Under 3D culture conditions, matrix properties can influence cellular stiffness^30^ because the extracellular cues are coupled to actin stress fibers through integrins and focal adhesion complexes to modulate the stiffness of cells.^52^ It is known that higher cell-mediated matrix degradability reduces matrix confinement and causes higher actomyosin contractility, leading to increased cellular mechanical stress. In agreement with this notion, our results show that increased matrix degradability is associated with increased cell stiffness. When the matrix is degradable by the cells, there is lower cell-cell contact and higher cell-matrix interactions, increasing cell contractility and cellular traction.^31, 53^ Higher cellular mechanical stress corresponds to the maintenance of stem cell biomarkers, and lower cellular mechanical stress promotes the cells to undergo differentiation.^53^ In agreement with this, an earlier study demonstrated the importance of matrix-mediated cell confinement and lower cell contractility in the formation of correctly polarized epithelial lumen.^54^ A recent study has demonstrated the relation between traction forces and cell stiffness in breast epithelial cells, where increased traction forces are accompanied by higher cell stiffness.^55^

Y27632 has been used previously to relieve cellular mechanical stress caused by actomyosin contractility, thereby decreasing cell stiffness.^31, 56^ In agreement with this notion, our results show that ROCK inhibition reduced cell stiffness. Inhibition of ROCK in primary acinar cells has decreased YAP transcriptional targets and prevented YAP nuclear localization, thus inhibiting the activation of mechanotransduction pathways.^45^ In 100DEG cultures, ROCK inhibition led to the loss of epithelial spheroid morphology, ectopic deposition of basement membrane proteins, and limited acinar differentiation. Without Y27632, 0DEG cultures expressed lower levels of stem/progenitor markers than the 50DEG cultures. Although comparable amounts of polarized structures were developed in 0DEG and 50DEG cultures upon Y27632 treatment, the stimulatory effects on acinar differentiation are more pronounced in the latter conditions. Thus, for the development of acini-like polarized structures, some degree of matrix confinement is needed.

Collectively, this study, for the first time, identified 3D culture conditions to achieve correct apicobasal polarization in hS/PC-derived multicellular assemblies that structurally and phenotypically resemble the native acinus. Upon neurotransmitter stimulation, luminal secretion of amylase granules was observed. We highlight the importance of maintaining optimal matrix confinement, cell-mediated matrix remodeling, and cellular stiffness to promote the formation of polarized lumen and stimulate acinar differentiation. This work represents significant progress in the development of bioengineered salivary gland mimetics for the treatment of radiation-induced xerostomia.

## CONCLUSION

Primary human salivary stem/progenitor cells were encapsulated in RGD-containing, mechanically matched HA hydrogels with varying percentages of MMP-cleavable crosslinks. The resultant cellular constructs were maintained in HepatoStim media with or without the ROCK inhibitor Y27632. Encapsulated single hS/PCs grew into multicellular spheroids by day 15. At the single-cell level, we discovered that increased matrix degradability correlated with increased cell stiffness and treatment with the ROCK inhibitor rendered cells softer. In non-degradable (0DEG) and 50% degradable (50DEG) matrices, ROCK inhibition led to the establishment of multicellular structures that were correctly polarized, as evidenced by the apical localization of GM130, apical-lateral localization of tight junction proteins, basal localization of β1 integrin and basement membrane proteins. and secretion of α-amylase into the lumen upon neurotransmitter stimulation. ROCK inhibition in 50DEG resulted in increased expression of acinar markers at the mRNA (*AQP5*, *SLC12A2*) and the protein (NKCC1, AQP5) levels with a concomitant decrease in the expression of ductal markers (*TFCP2L1*, *KRT7*). We conclude that an optimal level of cell stiffness and matrix degradability is desirable to guide the assembly of hS/PCs into multicellular spheroids with correct apicobasal polarization and acinar phenotype. This finding represents a significant leap forward in the field of salivary gland tissue engineering, paving the way for future advancements and potential clinical applications.

## ACKNOWLEDGEMENT

This work was supported in part by the National Institutes of Health (NIDCR R01 DE029655, NIDCD, R01DC014461) and National Science Foundation (NSF, DMR 1809612). The authors also acknowledge the use of facilities and instrumentation supported by NSF through the University of Delaware Materials Research Science and Engineering Center (DMR-2011824). Microscopy access was supported by grants from the NIH-NIGMS (P20 GM103446), the NSF (IIA-1301765), and the State of Delaware. This research also benefitted from the BioStore data management resource at the University of Delaware Bioinformatics Data Science Core (RRID: SCR_017696) supported by an NIH shared instrumentation grant (NIH S10 OD028725) and Delaware INBRE (P20 GM103446). We thank Dr. Jeffrey Caplan and Dr. Sylvain Le Marchand for their expert assistance in confocal imaging and image analysis. We thank Sanofi/Genzyme for generously providing HA.

