## Supplementary Information for "Promoting Polarization and Differentiation of Primary Human Salivary Gland Stem/Progenitor Cells in Protease-Degradable Hydrogels via ROCK Inhibition"

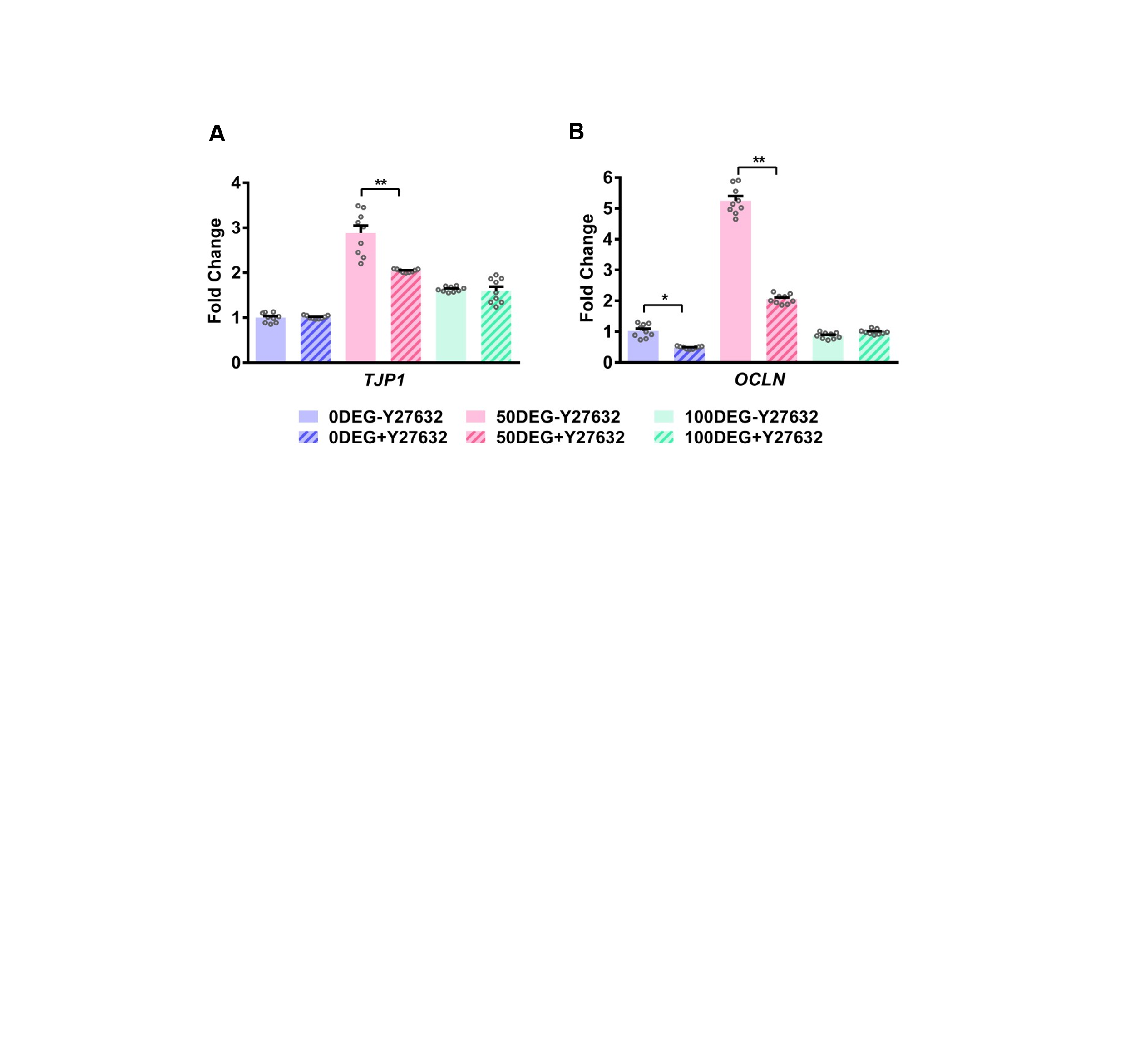

**Figure S1.** RT-qPCR analyses of day 15 constructs cultivated with or without Y27632 for the expression of tight junction proteins TJP1 **(A)** and OCLN **(B)**. Expression was normalized to Y27632-free 0DEG cultures. Error bars represent SEM. One-way ANOVA post hoc Tukey’s test was performed; ** indicates p < 0.01.

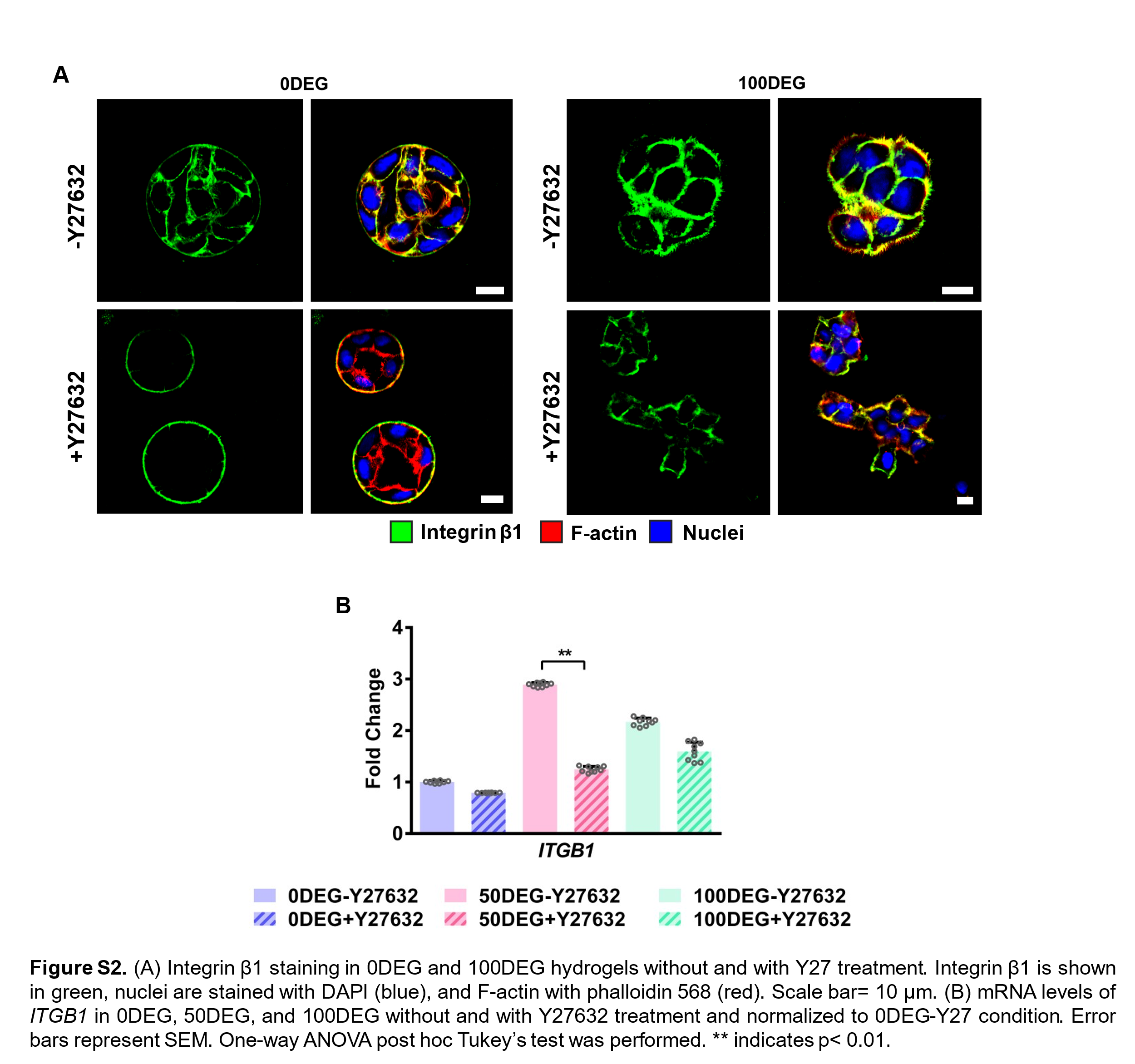

**Figure** **S2.** **(A)** Confocal images of day 15 0DEG and 100DEG constructs cultured with or without Y27632 and stained for Integrin β1 (green). F-actin and nuclei are shown in red and blue, respectively. Scale bar: 10 µm. **(B)** RT-qPCR analyses of day 15 constructs cultivated with or without Y27632 for the expression of ITGB1. Expression was normalized to Y27632-free 0DEG cultures. Error bars represent SEM. One-way ANOVA post hoc Tukey’s test was performed. ** indicates p < 0.01.

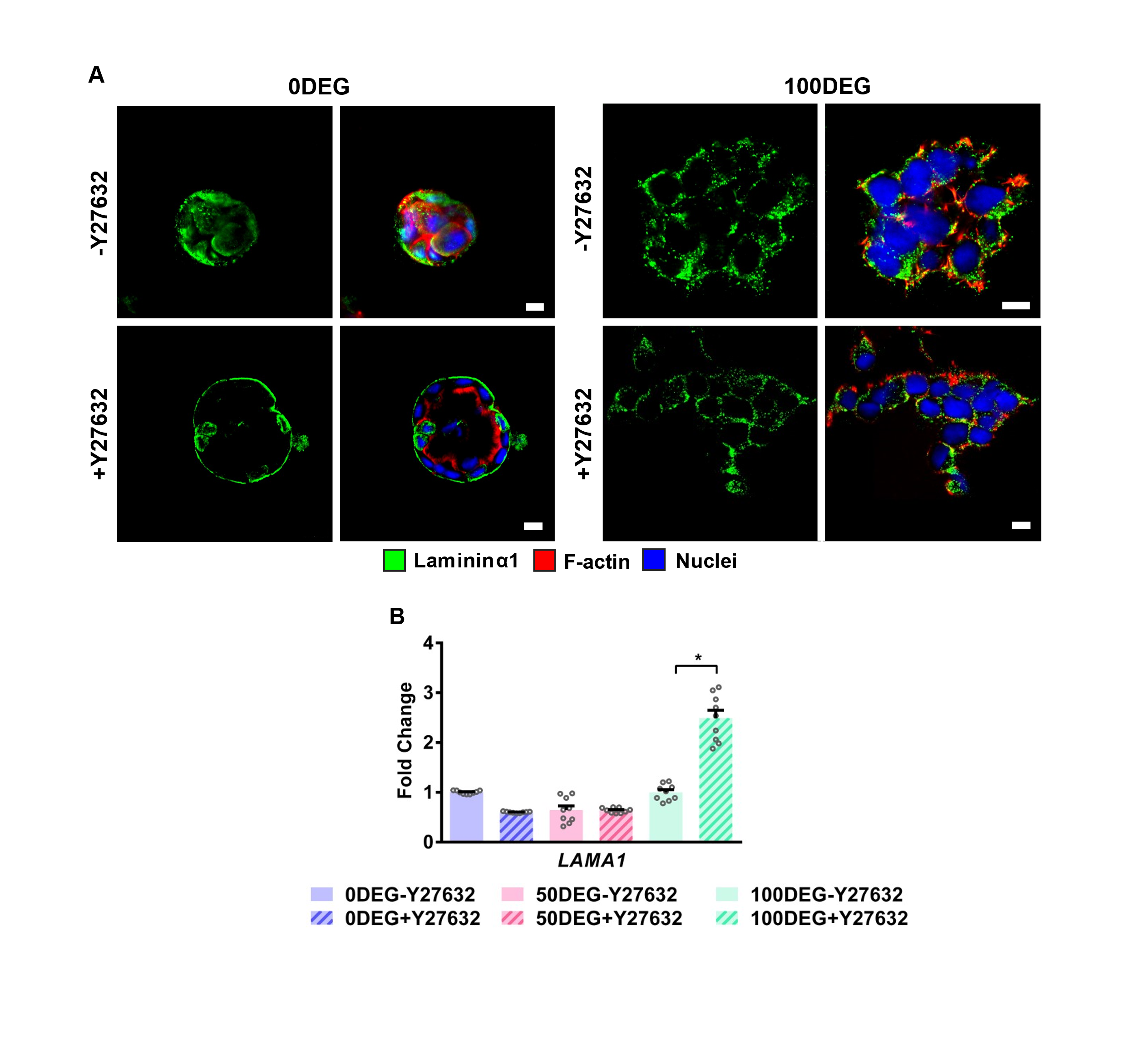

**Figure** **S3.** **(A)** Confocal images of day 15 0DEG and 100DEG constructs cultured with or without Y27632 and stained for laminin α1 (green). F-actin and nuclei are shown in red and blue, respectively. Scale bar: 10 µm. **(B)** RT-qPCR analyses of day 15 constructs cultivated with or without Y27632 for the expression of LAMA1. Expression was normalized to Y27632-free 0DEG cultures. Error bars represent SEM. One-way ANOVA post hoc Tukey’s test was performed. * Indicates p < 0.05.

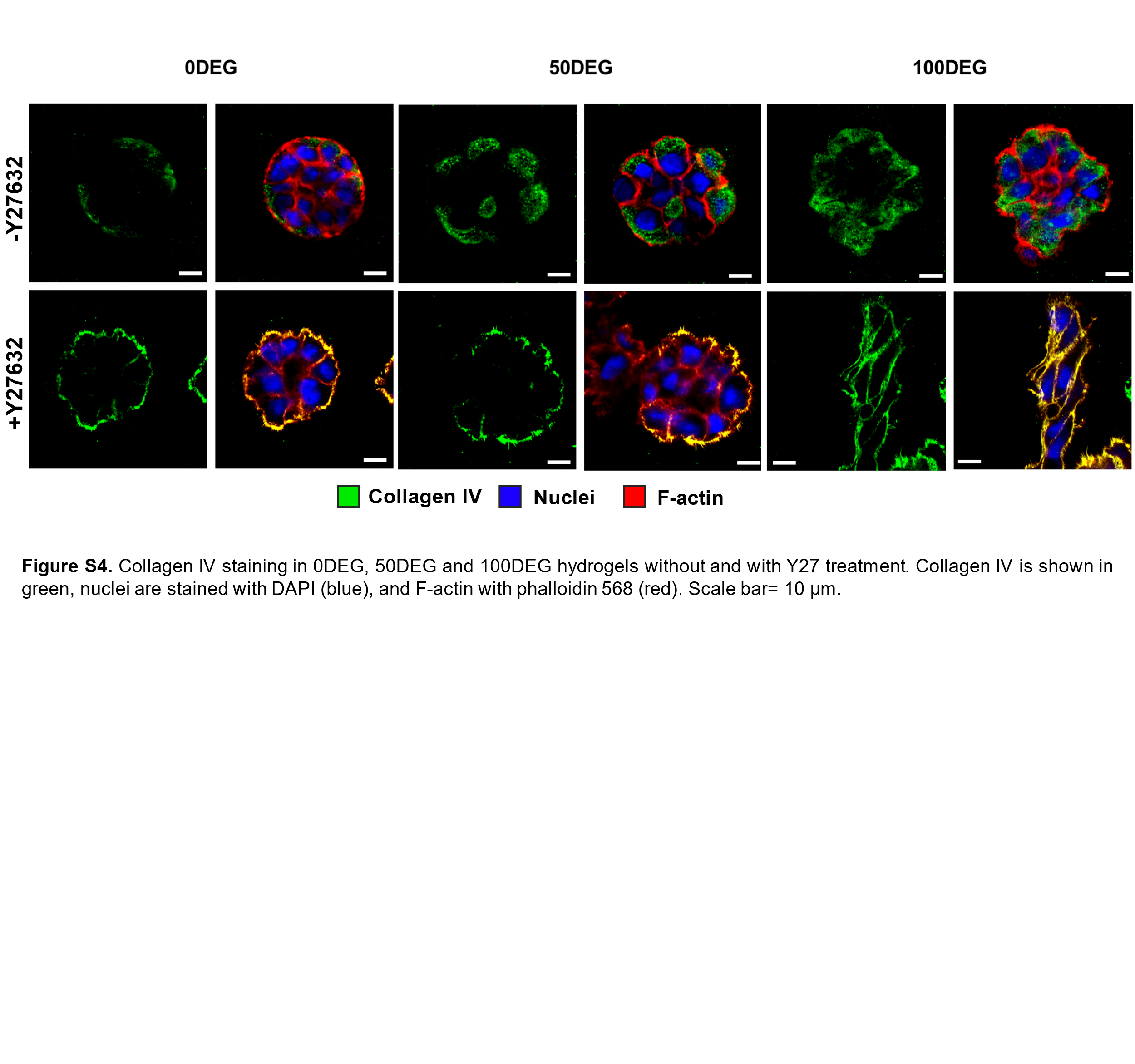

**Figure** **S4.** Confocal images of day 15 constructs cultured with or without Y27632 and stained for collagen IV (green). F-actin and nuclei are shown in red and blue, respectively. Scale bar: 10 µm.

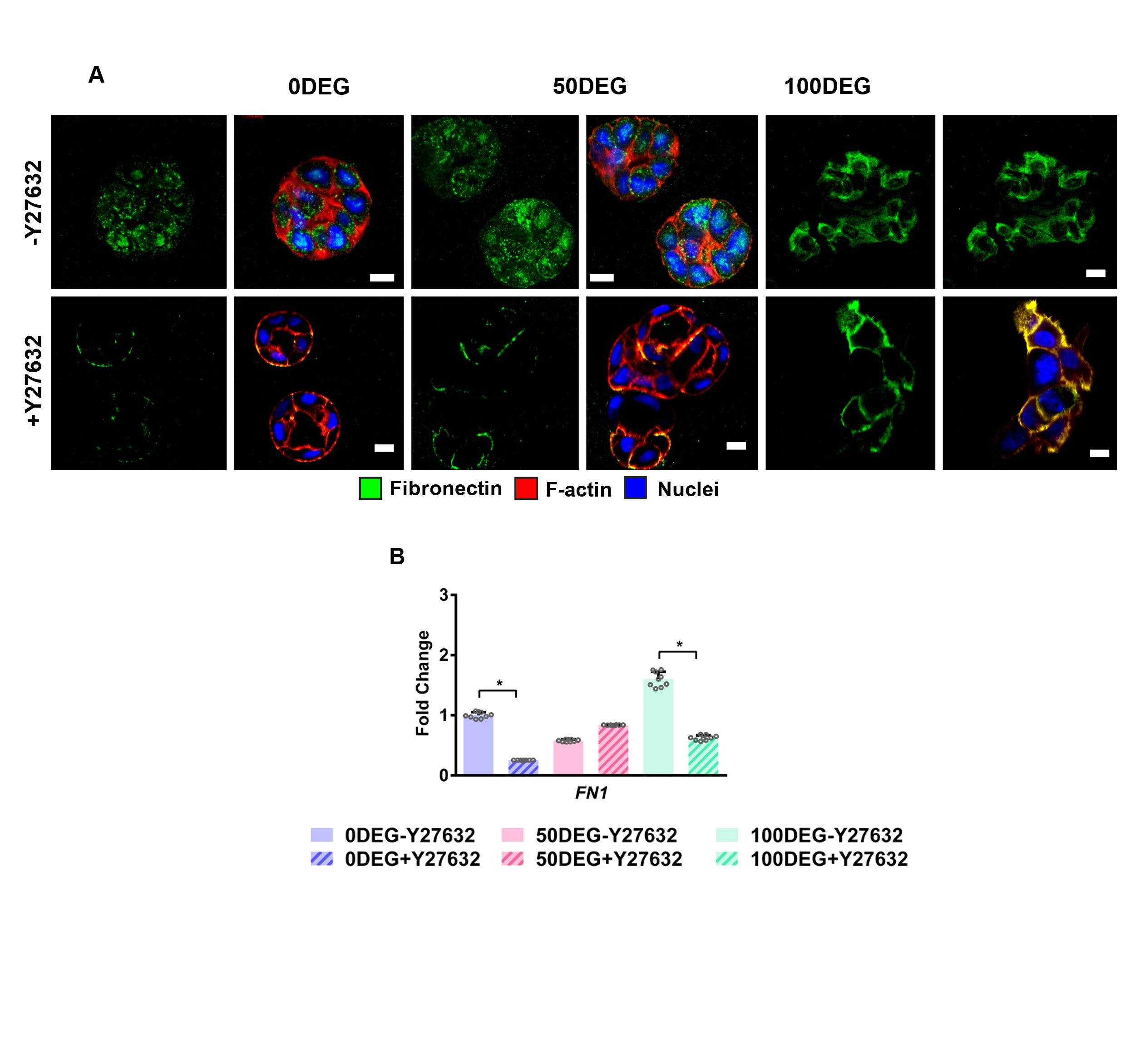

**Figure** **S5. (A)** Confocal images of day 15 constructs cultured with or without Y27632 and stained for fibronectin (green). F-actin and nuclei are shown in red and blue, respectively. Scale bar: 10 µm. **(B)** RT-qPCR analyses of day 15 constructs cultivated with or without Y27632 for the expression of FN1. Expression was normalized to Y27632-free 0DEG cultures. Error bars represent SEM. One-way ANOVA post hoc Tukey’s test was performed. * Indicates p < 0.05.

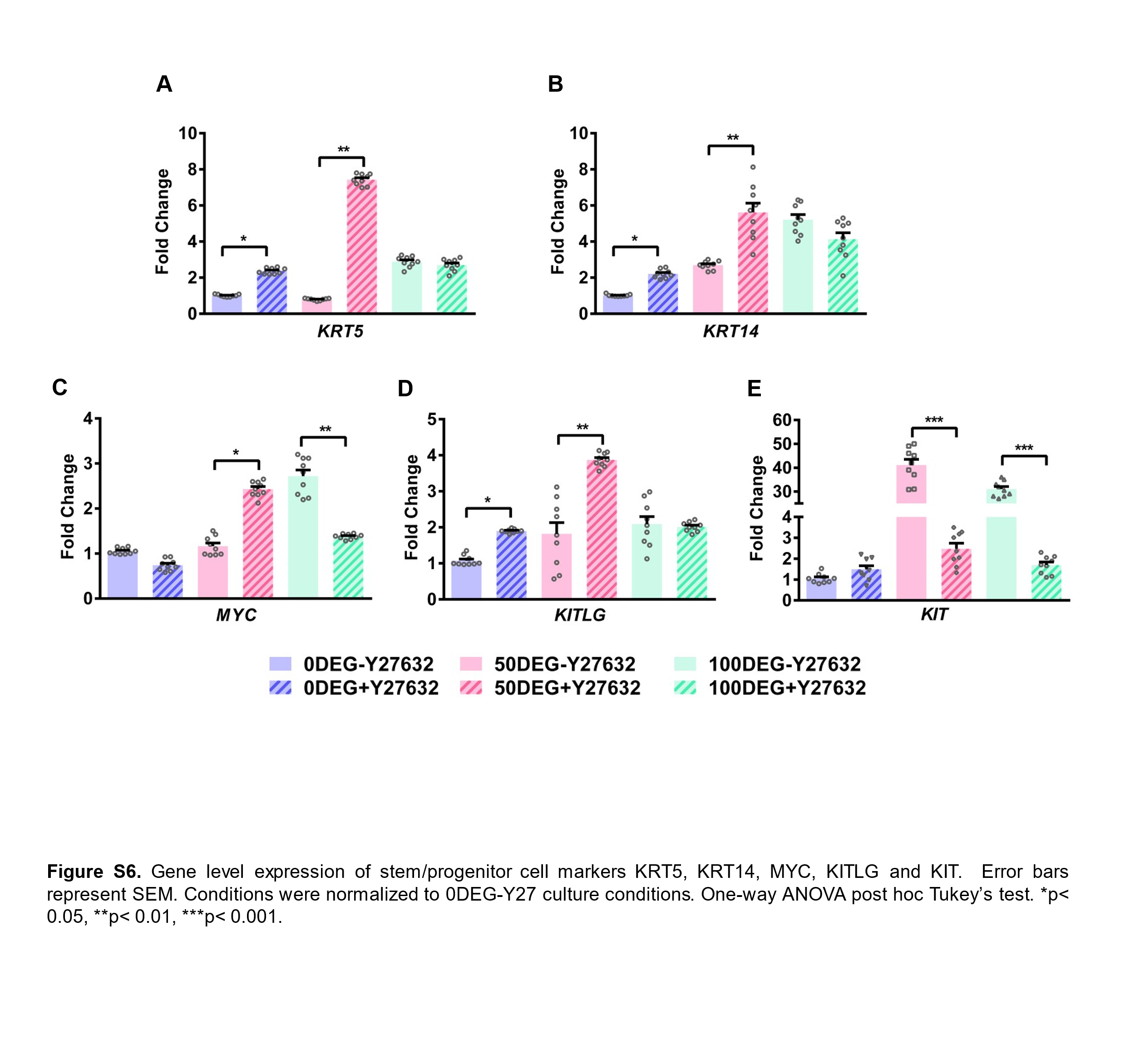

**Figure** **S6.** RT-qPCR analyses of day 15 constructs cultivated with or without Y27632 for the expression of progenitor (KRT5, KRT14) and stem cell (MYC, KITLG, KIT) markers. Expression was normalized to Y27632-free 0DEG cultures. Error bars represent SEM. One-way ANOVA post hoc Tukey’s test was performed. *, **, *** indicate p < 0.05, 0.01, and 0.001, respectively.

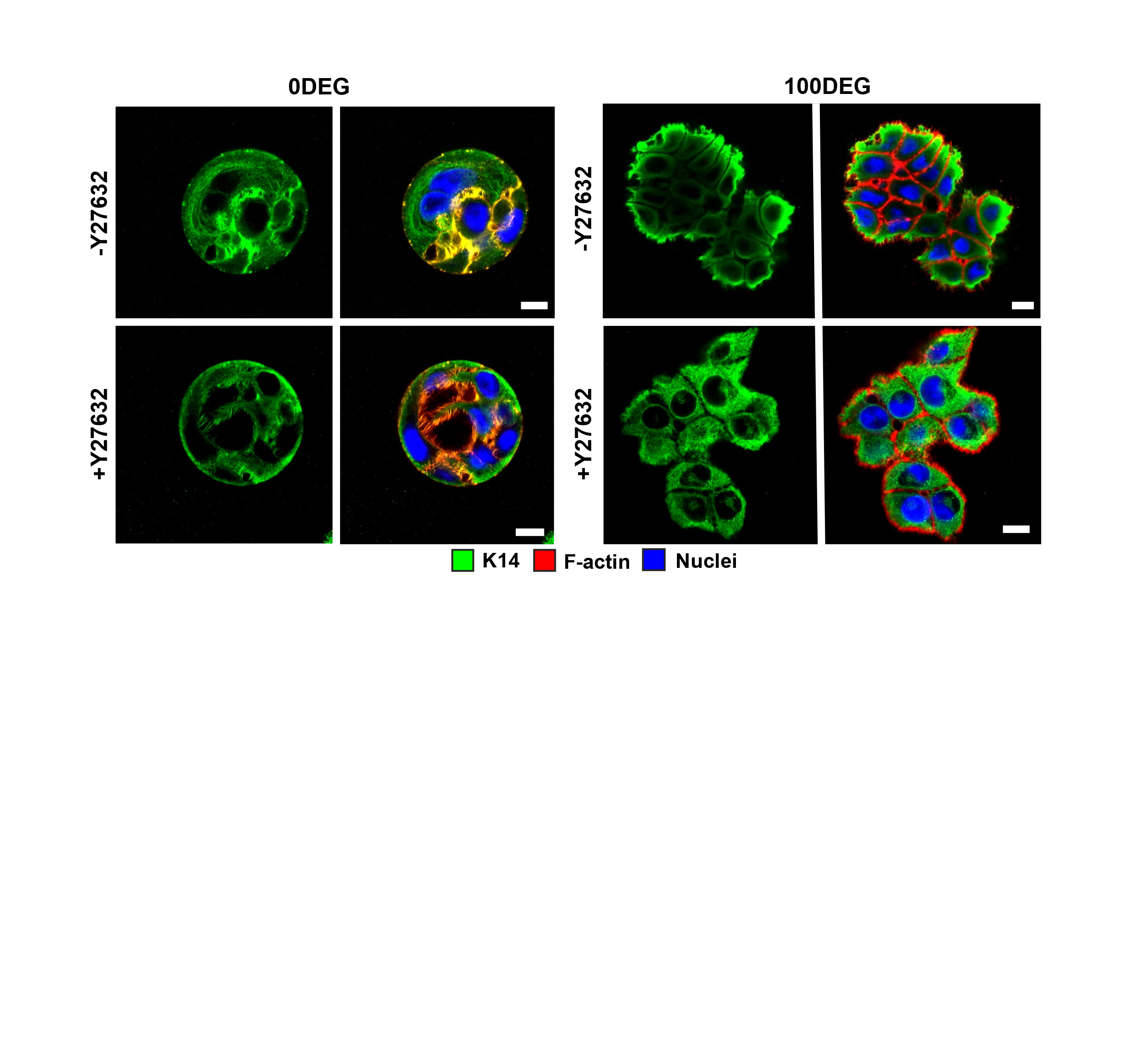

**Figure S7.** Confocal images of day 15 0DEG and 100DEG constructs cultured with or without Y27632 and stained for K14 (green). F-actin and nuclei are shown in red and blue, respectively. Scale bar: 10 µm.

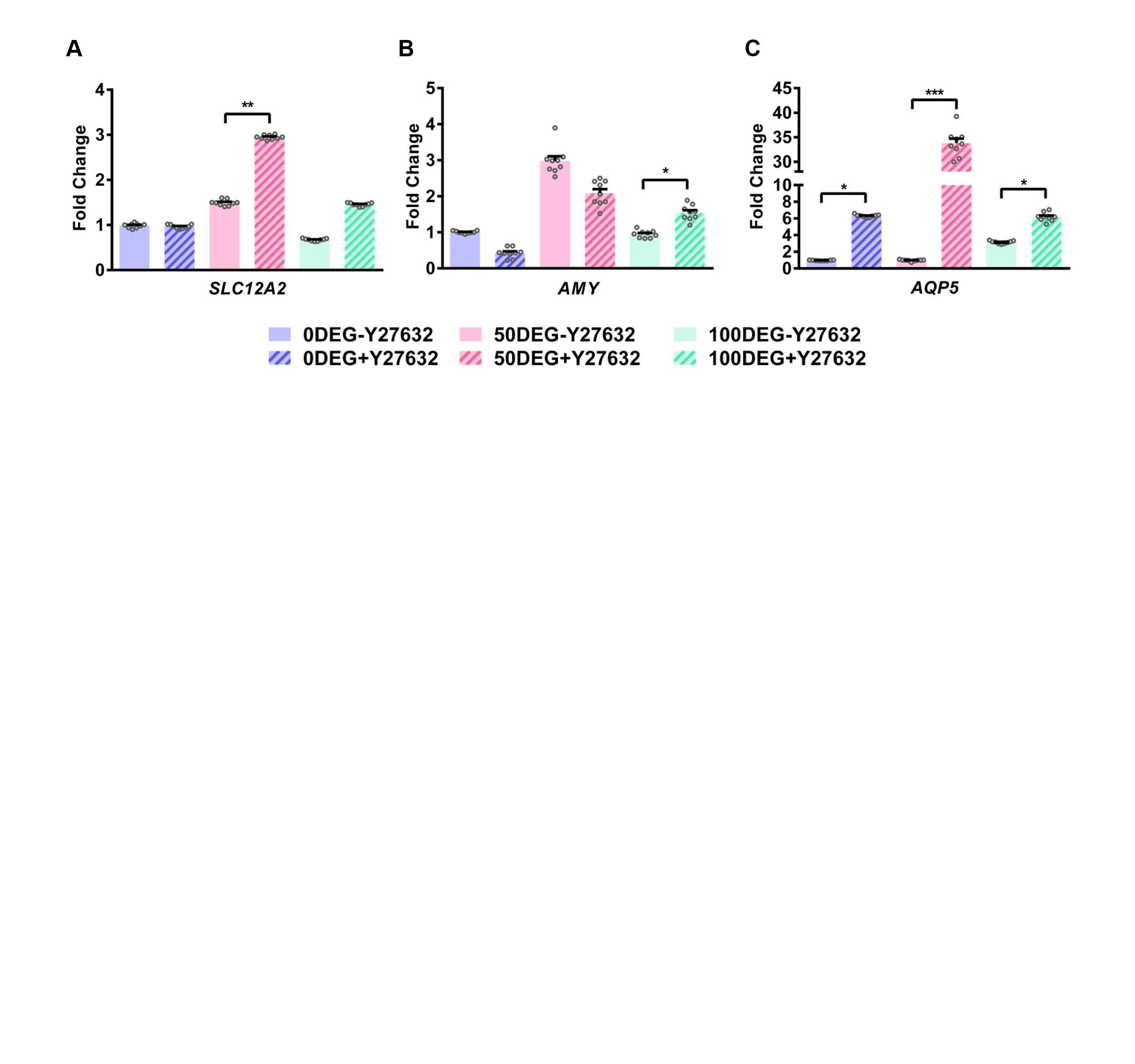

**Figure S8.** RT-qPCR analyses of day 15 constructs cultivated with or without Y27632 for the expression of acinar markers (SLC12A1, AMY, AQP5). Expression was normalized to Y27632-free 0DEG cultures. Error bars represent SEM. One-way ANOVA post hoc Tukey’s test was performed. *, **, and *** indicate p < 0.05, 0.01, and 0.005, respectively.

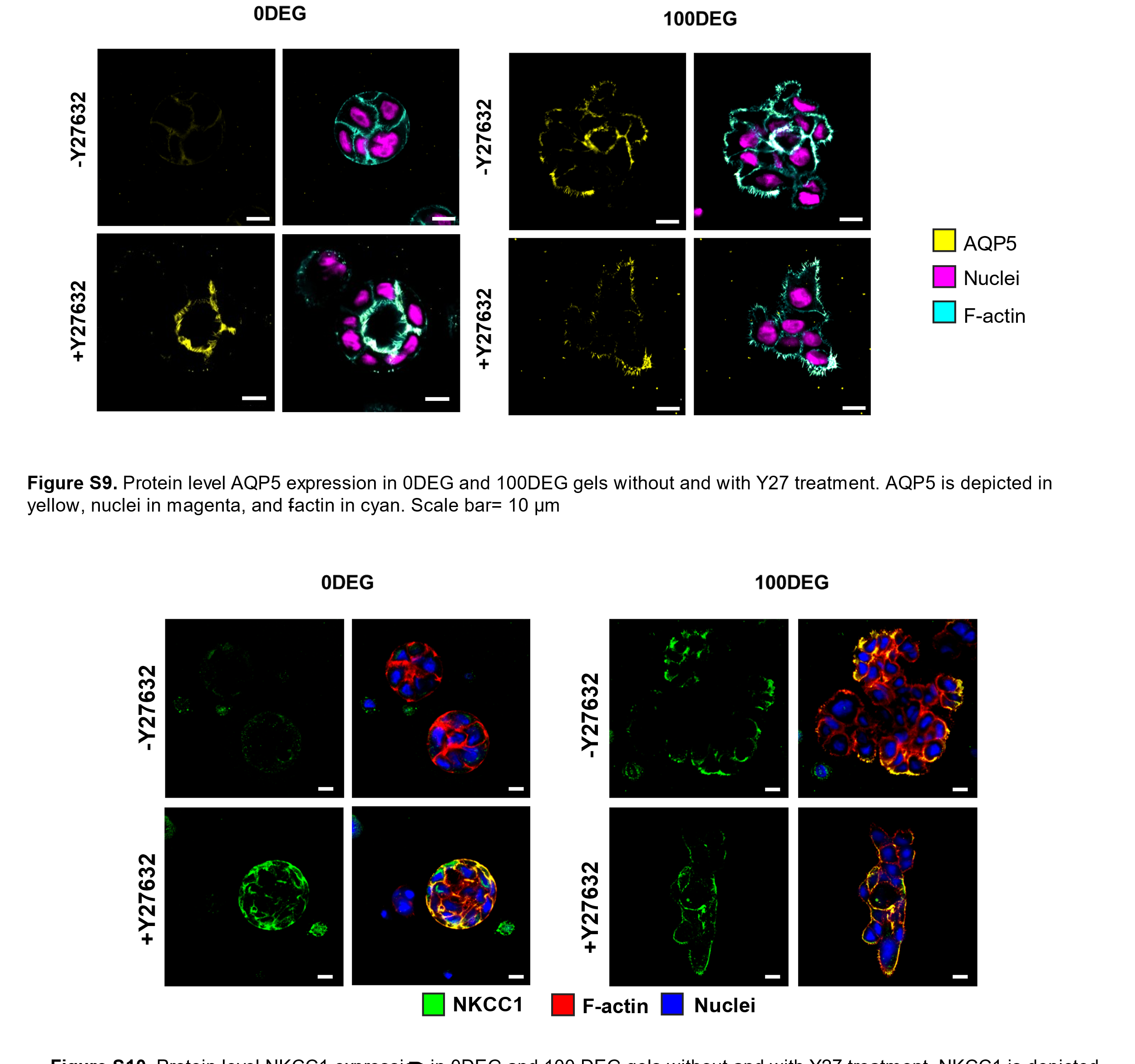

**Figure** **S9.** Confocal images of day 15 0DEG and 100DEG constructs cultured with or without Y27632 and stained for acinar marker NKCC1 (green). F-actin and nuclei are shown in red and blue, respectively. Scale bar: 10 µm.

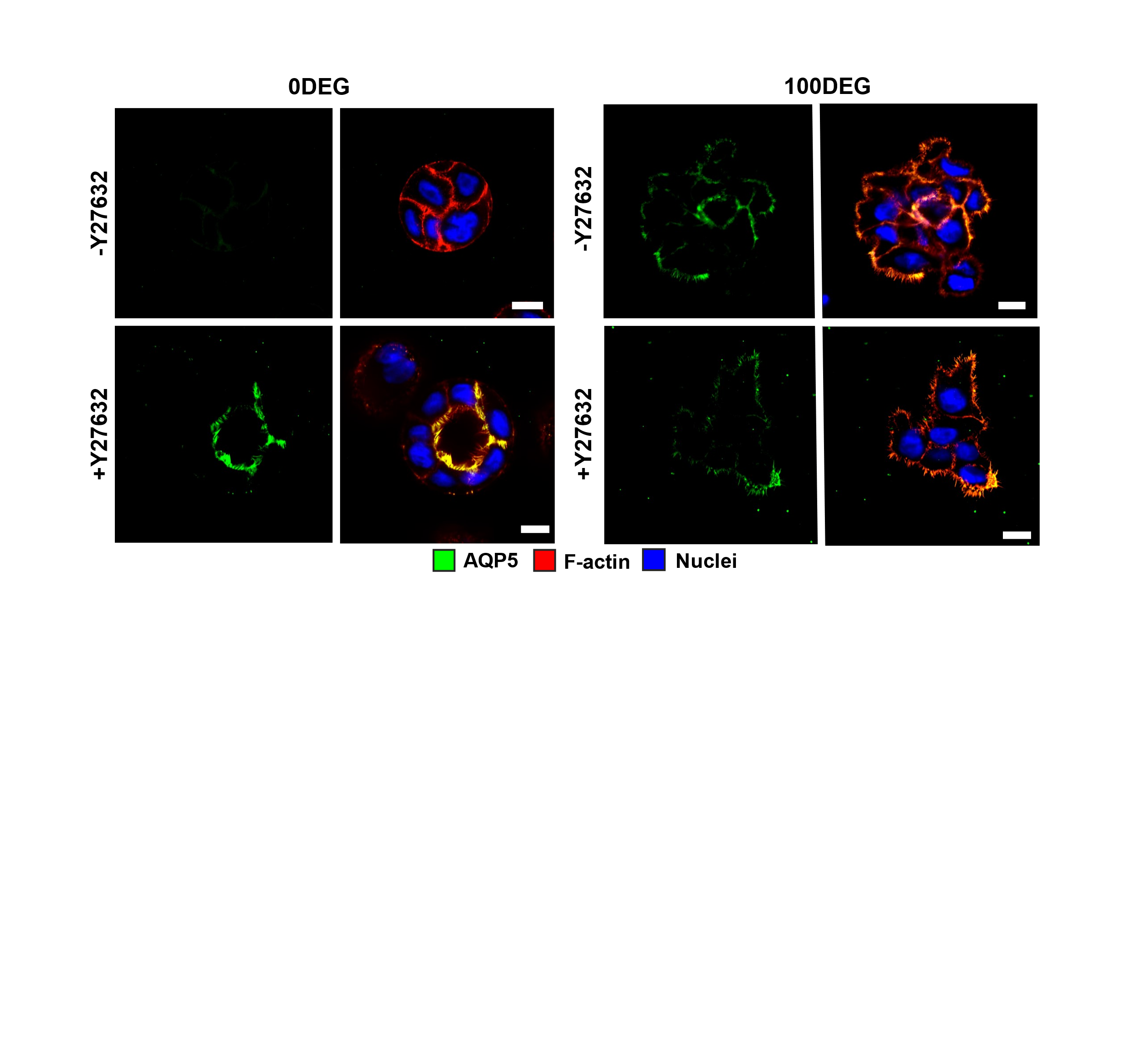

**Figure S10.** Confocal images of day 15 0DEG and 100DEG constructs cultured with or without Y27632 and stained for acinar marker AQP5 (green). F-actin and nuclei are shown in red and blue, respectively. Scale bar: 10 µm.

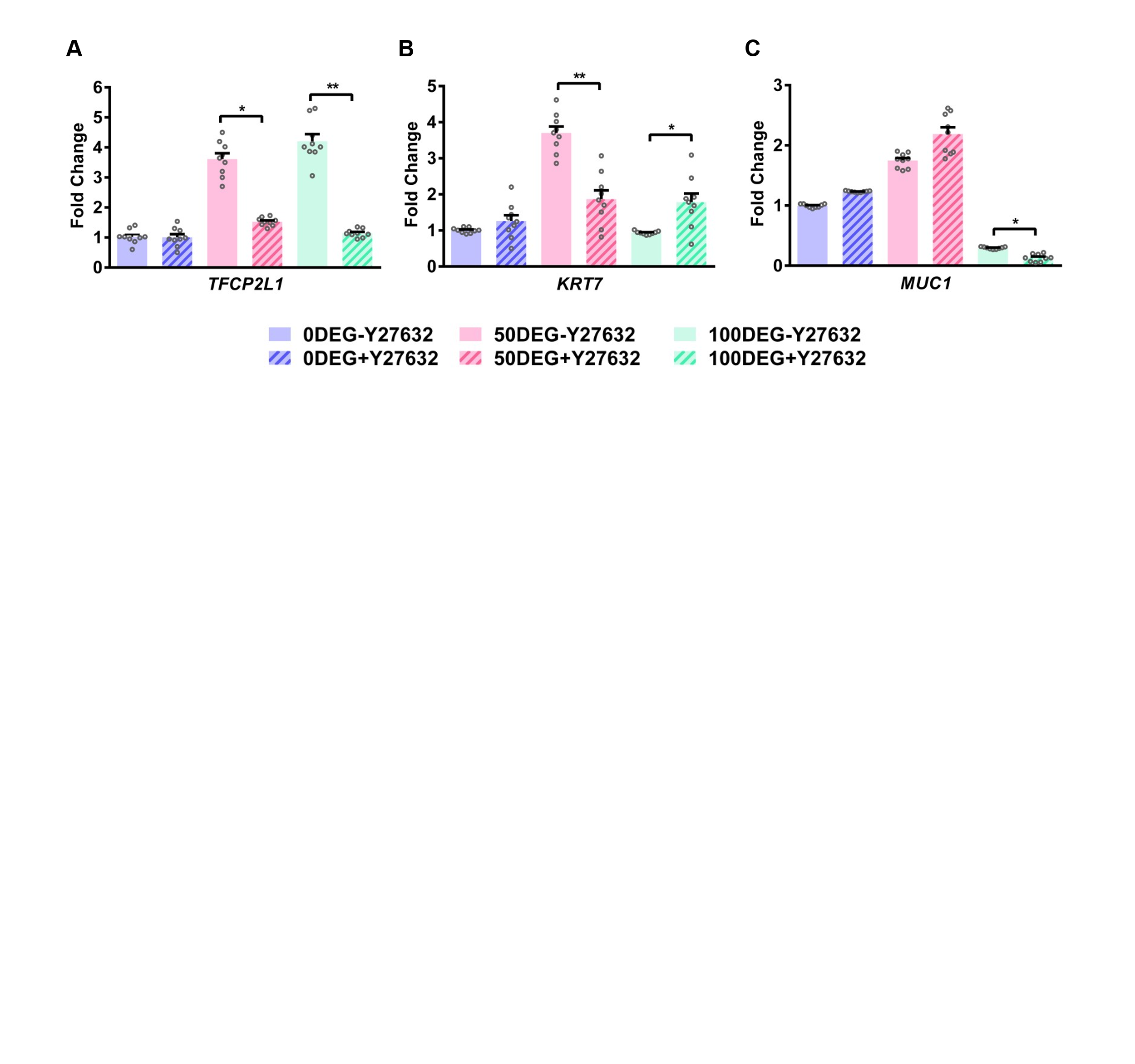

**Figure** **S11.** RT-qPCR analyses of day 15 constructs cultivated with or without Y27632 for the expression of ductal markers (TFCP2L1, KRT7, MUC1). Expression was normalized to Y27632-free 0DEG conditions. Error bars represent SEM. One-way ANOVA post hoc Tukey’s test was performed. * and ** indicate p < 0.05 and 0.01, respectively.

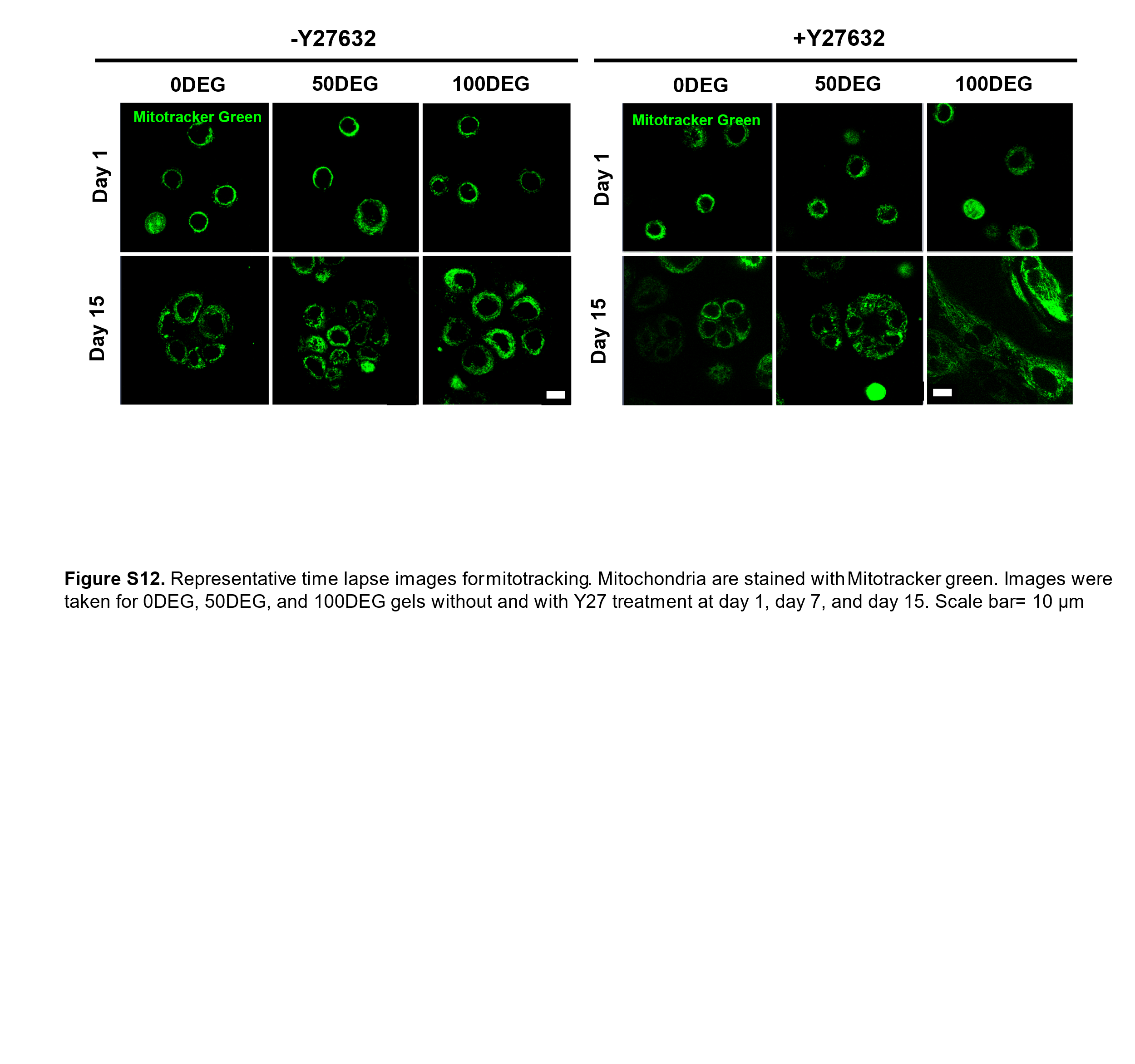

**Figure** **S12.**  Representative time-lapse images for particle tracking microrheology. hS/PCs were cultured in 0DEG, 50DEG and 100DEG with or without Y27632 for 1 and 15 days. Mitochondria was stained with mitotracker green, and its movement was tracked for 105 s.

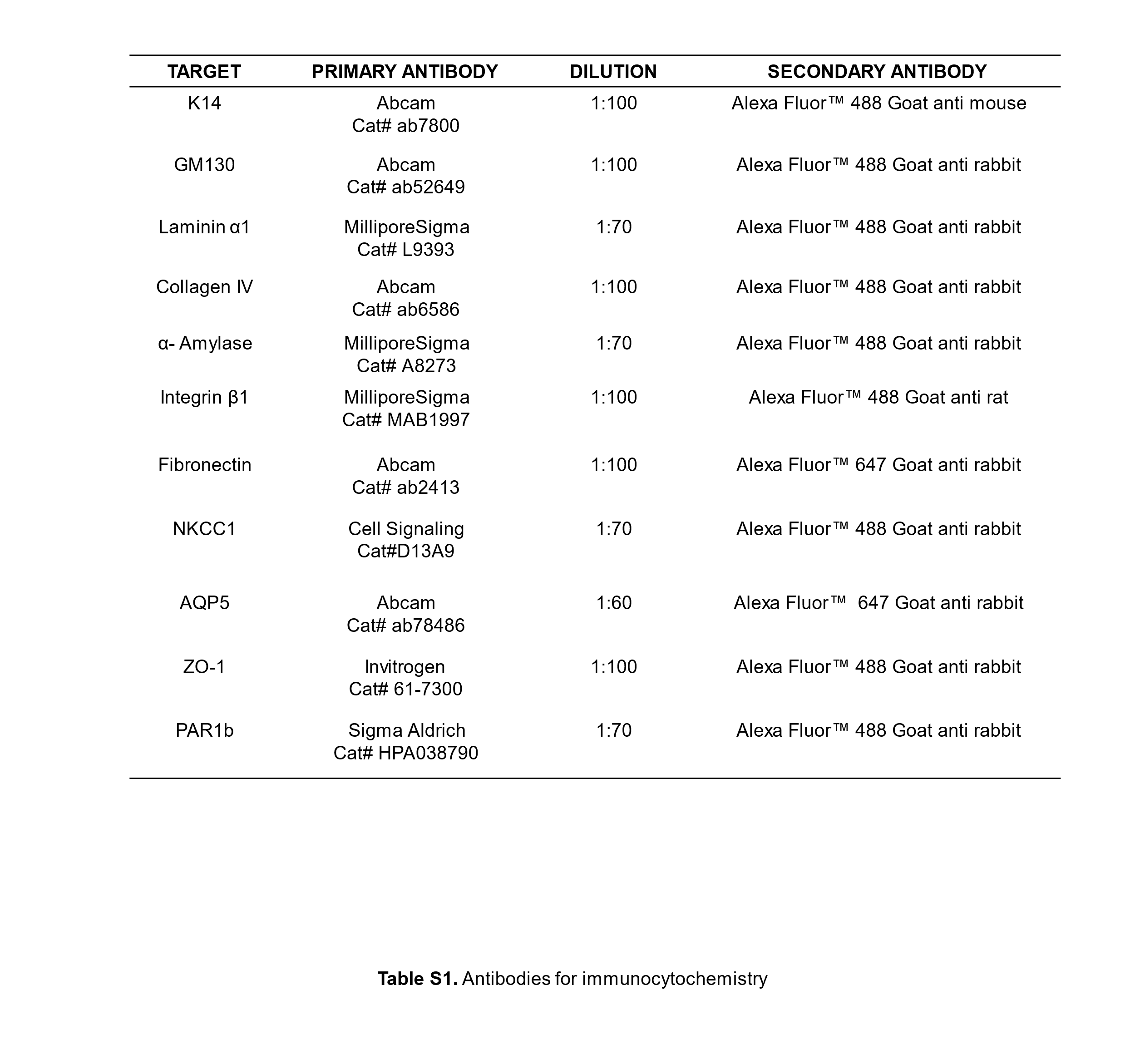

**Table S1.** Summary of antibody information for immunocytochemistry. The secondary antibodies were used at 1:200 dilution.

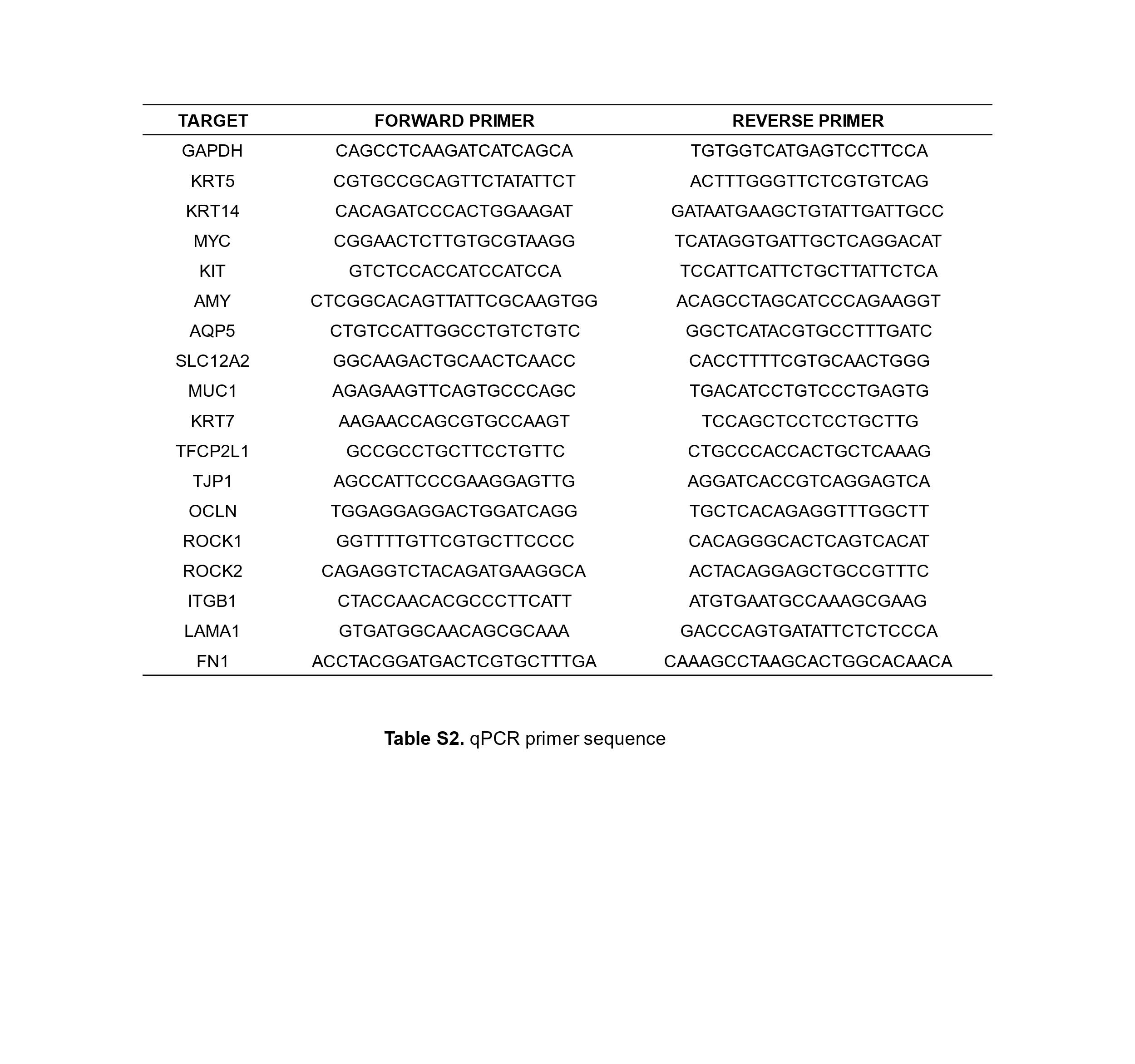

**Table S2.**  Summary of primers used in RT-qPCR analyses.

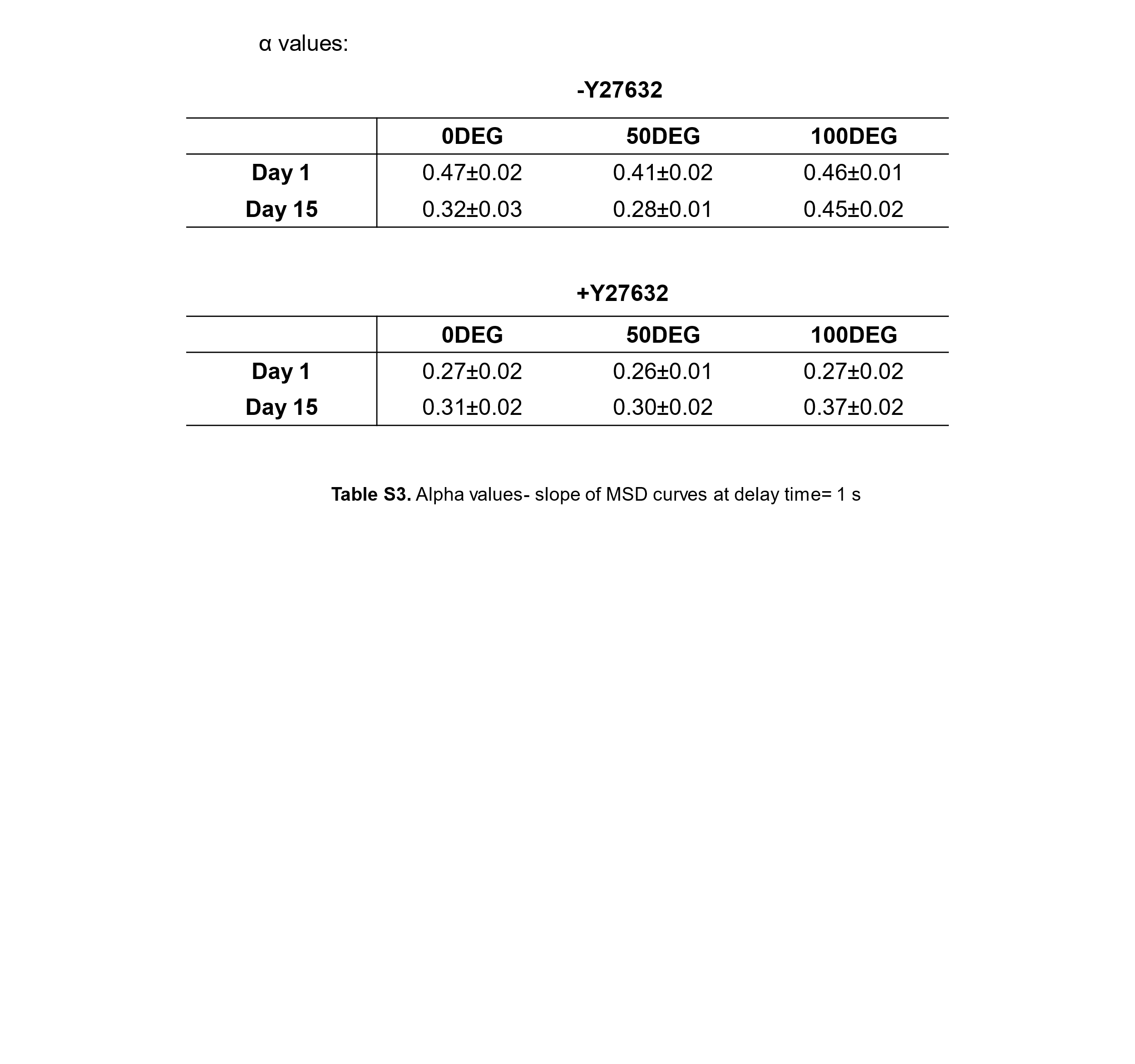

**Table S3.** Summary of α values calculated from the slope of MSD curves at delay time of 1 s. Error represents SEM.

**Supplementary Movie 1.** Confocal z-stack movie of hS/PC spheroids cultured in 0DEG gels without Y27632 and stained with F-actin (red) and nuclei (blue).

**Supplementary Movie 2.** Confocal z-stack movie of hS/PC spheroids cultured in 0DEG gels with Y27632 and stained with F-actin (red) and nuclei (blue).

**Supplementary Movie 3.** Confocal z-stack movie of hS/PC spheroids cultured in 50DEG gels without Y27632 and stained with F-actin (red) and nuclei (blue).

**Supplementary Movie 4.** Confocal z-stack movie of hS/PC spheroids cultured in 50DEG gels with Y27632 and stained with F-actin (red) and nuclei (blue).

**Supplementary Movie 5.** Confocal z-stack movie of hS/PC spheroids cultured in 100DEG gels without Y27632 and stained with F-actin (red) and nuclei (blue).

**Supplementary Movie 6.** Confocal z-stack movie of hS/PC spheroids cultured in 100DEG gels with Y27632 and stained with F-actin (red) and nuclei (blue).
